# Octopaminergic signaling contributes to thermal adaptation to elevation in African honey bees (*Apis mellifera*)

**DOI:** 10.64898/2026.05.10.724065

**Authors:** Florian Loidolt, Marco Mazzoni, Markus Thamm, Mark Otieno, Martin Hasselmann, Ricarda Scheiner

## Abstract

Adaptation to local environments enables species to thrive in diverse and challenging habitats. Steep elevational gradients provide a compelling natural adaptation laboratory, because abiotic conditions change progressively over short geographical differences. Given that elevation can strongly reshape physiology and behavior of insects, neuromodulatory systems offer a promising lens through which to examine elevation-specific adaptation. We challenged the hypothesis that adaptation to elevation involves octopaminergic signaling in honey bees (*Apis mellifera*), an important pollinator species occupying different elevations along East African mountains. We collected foragers from two distinct elevations at Mount Kenya (1,150 m and 1,900 m above sea level) and analyzed elevation-dependent changes in octopaminergic signaling. Tissue-specific analysis revealed a striking upregulation of all three octopamine β receptor genes in the thoracic flight muscles and elevated octopamine brain concentrations at high elevation. Expression differences in the brain and fat body were rather modest. We subjected CRISPR/Cas9-mediated octopamine β2 receptor knockouts to cold stress to study the function of octopaminergic signaling in thermoregulation. Loss of *AmOARβ2* reduced both the slope and amplitude of heating phases, indicating altered thermogenic dynamics. Together, these results identify the octopaminergic system as a central neuromodulatory regulator of thermogenic performance across elevations in honey bees. More broadly, our study highlights how modulation of conserved aminergic signaling pathways can shape physiological resilience to environmental gradients, pointing to a general mechanism by which insects adapt to changing thermal landscapes.

**Highlights:**

- Bees from high and low elevation differ in expression of octopamine β receptor genes and octopamine brain concentrations
- CRISPR/Cas9-mediated octopamine receptor knockout alters thermogenic behavior
- Octopaminergic signaling emerges as a key neuromodulator in thermal adaptation to elevation in honey bees

**Significance statement:** Animals living along mountain gradients must cope with rapidly changing temperatures, yet the mechanisms enabling this adaptation remain poorly understood. We show that honey bees from higher elevations have increased brain octopamine levels and enhanced expression of octopamine receptors in heat-producing flight muscles. Using gene editing, we demonstrate that disrupting one key receptor alters how bees generate heat under cold stress. These findings identify octopamine signaling as a central regulator of thermogenesis and reveal a mechanism by which insects adjust to colder environments. More broadly, our results highlight how conserved neuromodulatory systems can fine-tune physiological performance, offering insight into how insects may respond to changing climates and expanding environmental extremes.

## Introduction

The Western honey bee (*Apis mellifera*) occurs naturally across a wide range of altitudes, offering an excellent insect model for studying adaptation to elevation ^1,2^. The global distribution of this species demonstrates its remarkable capacity to thrive in diverse climatic conditions. Originating in Asia, *A. mellifera* has displayed an extensive adaptive radiation during multiple expansion events across Western Asia, Africa and Europe ^3^. The more than 30 described subspecies thrive in a variety of habitats, from high-precipitation tropical forests to hot desert or cold and harsh conditions at high latitude ^3,4^, requiring remarkable adaptability. A key strategy for temperature management is the thermoregulatory behavior in *A. mellifera* which accounts for its adaptive success even in high elevations ^5,6^. Unlike most insects, honey bees are capable of creating heat with their wing muscles. The increase of the thorax temperature is important for flight initiation and foraging at low temperatures but also for maintaining a constant hive temperature of around 34 °C to ensure optimal brood development and to survive cold conditions at high elevation ^7^.

Neuromodulation is an integral part of honey bee physiology and behavior. Recent research suggests a pivotal role of the neurotransmitter, hormone and neuromodulator octopamine ^8,9^ in honey bee thermogenesis ^10,11^. Octopamine is broadly expressed in neuronal and non-neuronal tissues and has conserved functions across invertebrates ^12,13^. To date, five octopamine receptor encoding genes are described for *Apis mellifera*: three β-adrenergic receptor genes *AmOARβ1, AmOARβ2, AmOARβ3/4*, and two α-adrenergic receptor genes *AmOARα1* and *AmOARα2*. They encode G-protein-coupled receptors with seven transmembrane domains affecting intracellular cAMP concentrations, apart from *AmOARα1*, which is calcium-linked ^14,15^. Octopamine plays a central role in physiology ^11,13,16^, learning and memory ^17–19^ and stress responses ^5,20^.

The pronounced environmental variation across elevations suggests that octopaminergic signaling may be under adaptive pressure, consistent with its involvement in a wide range of physiological and behavioral processes.

*A. mellifera* inhabits several mountain systems in East Africa at elevations well above 4,000 m, including Mount Kenya. Despite clear phenotypic and behavioral differences between lowland and highland populations, their genomes remain strikingly similar, with approximately 98.5% shared ^21^. Key elements of high genomic differentiation are located on chromosomes 7 and 9 and have been identified as chromosomal inversions, harboring multiple genes, including the three octopamine β receptors. Those structural genetic variants seem to be carried by honey bees of distinct elevations, fostering divergence of highland and lowland populations.

The high genomic similarity between highland and lowland honey bee populations, together with the consistent gene flow between them, suggests highly adaptive bee populations. *Apis mellifera monticola*, the darker honey bee found at high elevations, and *A. mellifera scutellata*, the more yellow honey bee prevailing in the lowlands, are therefore considered regional ecotypes rather than true subspecies ^21,22^. A recent investigation of honey bees along a mountain cline in Western Uganda provides a first, broad indication that honey bee populations at different elevations might exhibit adaptive differential octopamine receptor gene expression ^23^. Based on this evidence, we study the role of octopaminergic signaling in East African honey bees across elevations in detail, focusing on different tissues.

Here, we investigated how the octopaminergic system contributes to environmental adaptation in honey bees. We compared populations across two elevations on Mt. Kenya by profiling tissue-specific expression of all five known octopamine receptor genes and quantifying brain octopamine concentrations. To move beyond correlation, we disrupted octopaminergic signaling by generating a knockout of the candidate receptor *AmOARβ2* and measured how this manipulation altered thermogenic responses under acute cold stress.

## Material & Methods

### Study area and honey bee sampling

Samples of *Apis mellifera* were collected on Mt. Kenya located in central Kenya (approx. - 0.150°, 37.317°) in February 2024 (Figure 1 A). We selected two distinct elevations: a lowland elevation at approximately 1,150 m above sea level (ASL) characterized by savannah habitat and agricultural area, and a highland elevation at ca. 1,900 m ASL, which was occupied by montane forest and agriculture. Thus, the sampling sites were separated by roughly 700 m altitudinal difference. Samples were collected in a standardized way from three honey bee hives of collaborating local beekeepers per elevation (Figure 1 A; location, colony coordinates provided in Supplementary Table S1). The entrances of the hives were sealed, and returning foragers were captured in glass vials followed by immediate freeze-killing with Presto cooling spray (European Aerosols GmbH, Haßmersheim, Germany). Bees were quickly transferred to small plastic screw-lid tubes and placed on dry ice to preserve RNA and biogenic amine state. We sampled five bees per colony at both elevations for octopamine receptor mRNA expression and an additional ten bees per hive for the biogenic amine brain concentrations. Samples were stored at -80 °C at the University of Embu research facility, shipped on dry ice to the University of Würzburg (Germany), and stored at -80 °C until further processing.

**Figure 1.**
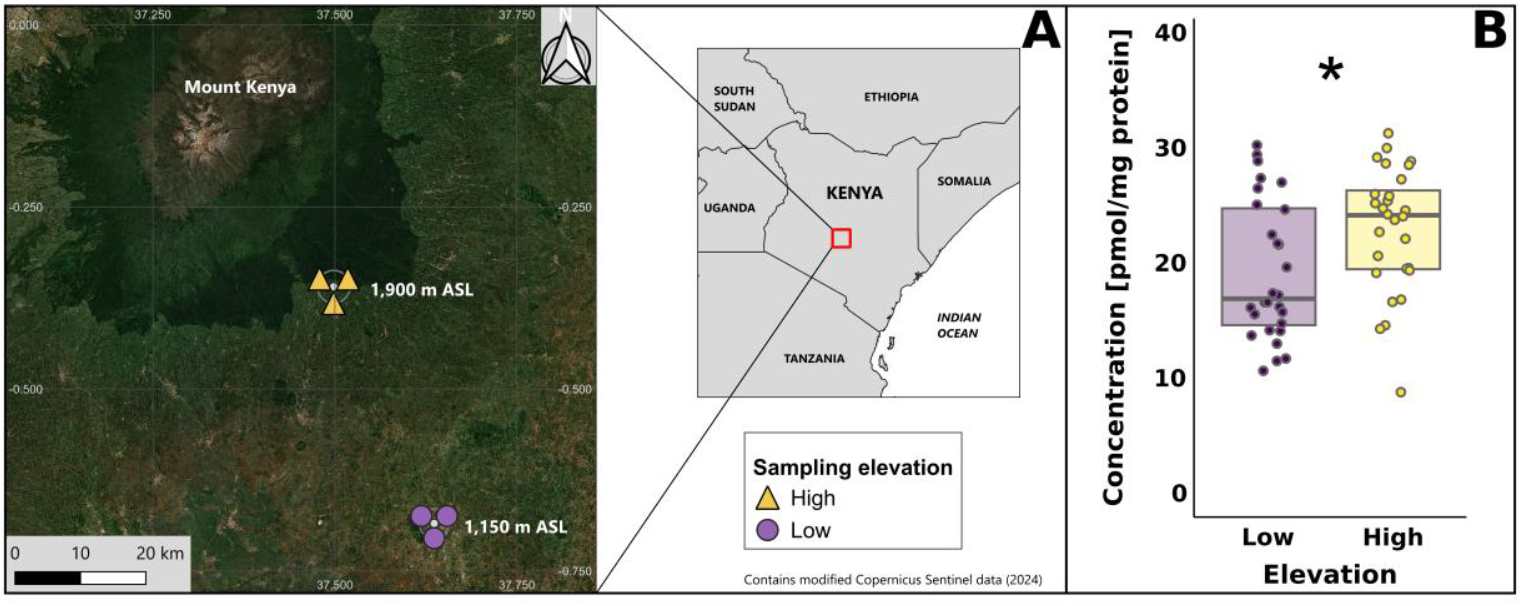
Study area and elevation-dependent brain octopamine concentration. **A**. Study area and sampling sites in Mt. Kenya region, Kenya. The map shows the location of the honey bee colonies for sampling at low elevation (purple circles, 1,150 m ASL, N = 3) and high elevation (yellow triangles, 1,900 m ASL, N = 3). Basemap: Sentinel-2 cloudless mosaic, with modified Copernicus Sentinel data from 2024 ^46^. Overview map: Natural Earth (2024). For detailed colony coordinates information see Supplementary Table S1. **B**. Octopamine concentration in the brain of honey bees sampled at low (purple) and high elevation (yellow). N = 28 each, significant differences indicated by asterisks: * = *P* < 0.05. For detailed statistics see Table 1.

### Gene expression and quantification of octopamine concentration in the brain

For a high-resolution gene expression of all octopamine receptor genes, we analyzed brain, thorax muscle and fat body tissues separately. After sampling, bee heads were placed in RNAlater-ICE (Thermo Fisher Scientific, Waltham, USA). Each head was placed into a tube with 200 µl of -20 °C pre-cooled RNAlater-ICE, while the abdomen was cut off and placed into 500 µl of the same solution. For proper penetration of the tissue, antennae were removed and several holes were cut into the head capsule. The tubes were placed at -20 °C for seven days and dissected at room temperature. The head capsule was opened, and the total brain was dissected in liquid nitrogen. The abdomen was carefully opened with scissors to take out the gut and stinger apparatus first. After removal of the tracheal tissue and the abdominal ganglion chain, the exoskeleton with the attached fat body tissue remained. Thorax muscle dissection was performed under liquid nitrogen. The cuticle and non-muscle tissue were carefully scratched off with two fine pincers before the entire thorax muscle was extracted. Total RNA was extracted from each tissue using the GenUP Total RNA Kit (biotechrabbit, Henningsdorf, Germany). In addition to the protocol, we performed a 15 min DNase digestion step at room temperature on the column-bound RNA to remove DNA from the samples (30 µl DNase I, 5 µl 10 x Reaction Buffer, 15 µl H_2_O; Lucigen Corporation, Middleton, USA). Final RNA elution was performed in 80 µl H_2_O for the brain and 100 µl for fat body and thorax muscle. cDNA was synthesized with the Biozym cDNA Synthesis Kit (Biozym, Hessisch Oldendorf, Germany), using standardized 200 ng total RNA per sample and tissue. Gene expression was analyzed using quantitative real-time PCR (qPCR) in triplicate per biological sample and tissue type. A tissue-specific calibrator cDNA was added to every qPCR run for calibration across runs. Every PCR reaction had a total volume of 20 µl (10 µl 2 x qPCR S’Green BlueMix (Biozym, Hessisch Oldendorf, Germany), 5 µl cDNA, 4.2 µl H_2_O, 0.4 µl forward primer, 0.4 µl reverse primer) and was run at a Rotor Gene Q (QIAGEN, Hilden, Germany) with these temperatures: initial denaturation for 2 min at 95 °C followed by 40 cycles of 1) denaturation for 5 sec at 95 °C and 2) primer annealing and elongation for 30 sec at 60 °C, final extension for 90 sec at 60 °C, and ultimately a melting ramp from 60 °C to 95 °C with an increase of 1 °C every 5 seconds. The expression levels of the five target genes (*AmOARα1, AmOARα2, AmOARβ1, AmOARβ2, AmOARβ3/4*) were quantified relative to the expression of three stable reference genes (*AmRpL10, AmRpL19, AmRpL32*; for oligonucleotide sequences and gene IDs see Supplementary Table S2) with the qBase algorithm ^24^. Sample triplicates were checked for consistency in their cycle thresholds (ct) ^25^. In case of a replicate ct-value deviating more than 2 from the median, it was removed as a methodological outlier, leading to ct-value calculation from fewer replicates. If only one replicate per sample remained, a meaningful ct-value determination was not possible and the sample was excluded from the dataset of the respective gene.

The concentration of octopamine was analyzed in the brains of the bees ^8,26^. Brain dissection was performed under liquid nitrogen. Cuticle, muscle and tracheal tissues, hypopharyngeal glands and outer compound eye tissues were carefully removed with sharp forceps, and the brain including all neuropils was dissected and stored at -80 °C until further processing. High-performance liquid chromatography and Bradford protein measurement were performed according to established protocols ^11^. Frozen brains were placed into 60 µl of an extraction solution (10 pg/µl 3,4-dihydroxy-benzylamine (DHBA) as internal standard in 0.2 M perchloric acid). After centrifugation at 21,130 g and 4 °C for 2 min, an ultrasonic bath was used to break down the tissue (10 min in ice water). Samples were incubated in the absence of light on ice for 20 min and centrifuged at 21,130 g and 4 °C for 14 min. The supernatant containing the biogenic amines was collected and analyzed via HPLC-ESD, while the pellet was frozen at - 80 °C for the Bradford assay. 20 µl of sample volume were run at a 3 µm reverse phase column (BDS-Hypersil-C18, 150 x 3 mm, 130 Å pore size; Thermo Fisher Scientific, Waltham, USA) equipped with a two coulometric cell ECD-3000RS detector (6011RS ultra-analytical cell; Thermo Fisher Scientific, Waltham, USA). Run conditions comprised a special mobile phase (15% methanol (v/v), 15% acetonitrile (v/v), 85 mM sodium phosphate monobasic, 1.75 mM sodium dodecylsulfate, 0.5 mM sodium citrate, ultrapure water), a stable pH environment of 5.6 ± 0.01 guaranteed by phosphoric acid, a flow rate of 0.5 ml/min with a flow ramp of 0.02 ml/min^2^, and the two detector cells set to 425 mV (DHBA) and 800 mV (octopamine) working potential. Run data analysis was performed with the software Chromeleon v. 7.2.10 (Thermo Fisher Scientific, Waltham, USA) by peak integration and calibration. In addition to HPLC amine level detection, we performed Bradford protein quantification ^27^ which allows normalization of the octopamine concentration on the total protein amount considering inter-individual size variation. The stored pellets were first resuspended in 100 µl of 0.2 M NaOH and homogenized at 40 Hz for 3 min using a metal bead and a TissueLyser LT (QIAGEN, Hilden, Germany). After 15 min of incubation on ice, the samples were centrifuged (5 min, 9,391 g) and 10 µl of the supernatant was added to 1 ml ROTI-Nanoquant (Carl Roth, Karlsruhe, Germany). Bovine serum albumin (BSA) served as standard for calibration in the respective concentrations of 1, 2, 3, 5, 10, and 20 µg/ml BSA Fraction V (Carl Roth, Karlsruhe, Germany) in the 1 x ROTI-Nanoquant solution. The analysis of the samples and the calibrator in a 96-well plate was performed with an Infinite 200 Pro plate reader (Tecan, Männedorf, Switzerland).

### Knockout of *AmOARβ2*

*AmOARβ2* knockout experiments were performed at the University of Würzburg (Germany) in August and September 2023 to provide safety level 1 laboratory environment for genetic manipulation of organisms. Because CRISPR-mediated editing was not feasible in the local mountain honey bee populations, *A. m. carnica* was used instead, with the assumption that gene function is conserved at the subspecies level. The gene *AmOARβ2* (National Center for Biotechnology Information – NCBI: Gene ID 412896, https://www.ncbi.nlm.nih.gov/gene/412896) is encoded on chromosome 7 of the *A. mellifera* genome and comprises 11 annotated exons. Most of the 12 available transcript variants share identical early coding DNA sequences (CDS) and thus the N-terminal protein region. Hence, we chose XM_016913140.2 as one of the longest transcript variants (5,109 bp linear mRNA) for the selection of potential single guide RNA (sgRNA) targets. Putative crRNA target sites were identified using Benchling (https://www.benchling.com/). Selection criteria followed Değirmenci et al. ^28^: 20 bp length, NGG PAM, and a G base at the 5′-end. We opted for candidates with high on-target scores (> 50%) indicating efficient DNA double-strand cleavage, and high off-target specificity scores approaching 100% to avoid unwanted interactions in other regions of the genome. We verified exon localization to avoid targets spanning exon-intron junctions with the software Splign (https://www.ncbi.nlm.nih.gov/sutils/splign/splign.cgi). sgRNA secondary structure and accessibility of the 20 bp targeting region were assessed using the Vienna RNAfold server (http://rna.tbi.univie.ac.at//cgi-bin/RNAWebSuite/RNAfold.cgi). This tool also allowed us to predict the presence of the required Cas9-interaction loops. In addition, we decided that the targets should be at a rather early position (maximum at ∼400 bp) in the CDS with a total length of 1,314 bp to increase the probability of function-disrupting indels. From these filters we chose the three most promising crRNA candidates for empirical testing (Supplementary Table S3). Three sgRNAs were produced in-house as described in Değirmenci et al. ^28^ by overlapping Phusion PCR using crRNA- and tracrRNA-containing primers, followed by DNA cleanup and in vitro transcription with T7 polymerase (RiboMAX™ Large Scale RNA Production Systems; Promega, Madison, USA). The synthesized sgRNAs were purified, quantified and stored in aliquots at -80 °C. Pilot injections testing three concentrations of each sgRNA candidate assessed both the larval hatching rate and the mutation frequency; these metrics were combined into a composite mutant rate to identify the most effective treatment (Supplementary Table S3). The sgRNA selected for the main experiment, showing the highest mutant rate, targeted the positive (+) strand at CDS position 122 (5′-GAACGCGAGCTACTCGAGCG-3′) and was used at an sgRNA concentration of 92 ng/µl in the injection mix.

Ten *A. mellifera* colonies with naturally mated queens were maintained to provide freshly-laid eggs. Egg collection and handling, microinjection and *in vitro* rearing followed established protocols ^28,29^. Each egg was injected with 400 pl of the injection mixture (92 ng/µl sgRNA and 3.13 µM Cas9). Non-injected eggs served as wild-type controls. Egg injections for the experiment resulted in a survival rate of approx. 70% after 24 hours for each of the three replicate injection days (Supplementary Table S4). Larval hatching rate was similar throughout days (Supplementary Table S4). During larval and pupal stages, dead individuals were removed regularly to avert the spread of mold and infections. Pupae were inspected twice daily during the eclosion period, and freshly emerged workers were placed into small cages. Cages were maintained at 30 °C and 60% relative humidity. All bees could consume the provided food *ad libitum*, i.e. 50% sucrose solution, pollen paste made from thawed pellets with 70% invert sugar solution, and water.

Artificial rearing of honey bee workers in the laboratory from larval stage to imago did not differ between treatments in survival (sgRNA-injected vs. non-injected wild types: *P* = 0.85, Fisher’s exact test; Supplementary Table S5), implying that *AmOARβ2* mutations were not lethal. The same was true for the eclosed adult honey bees during the period of ageing until the age of 14 days, when the behavioral tests were performed (*P* = 0.53, Fisher’s exact test; Supplementary Table S6).

### Behavioral cold stress assay

The reared honey bees were maintained in cages until 14-days-old, when they were fully capable of heating ^30^. Honey bees were separated into small glass vials and tested in groups of six, randomly picked from both wild-type and sgRNA-injected treatment groups. The glass vials remained at room temperature for 15 min to help the bees acclimate. Subsequently, we carefully fed each bee with 50% sucrose solution *ad libitum* and incubated the vials on ice for 5 min to induce chill coma. The immobilized bees were transferred to individual petri dishes (3 cm diameter). The petri dishes had a mesh top cover allowing for clear thermographic recording from above and ventilation while the bees could move freely.

The behavioral cold stress assay was performed in a Binder MK 56 incubator (Binder, Tuttlingen, Germany). The temperature was set to a constant 18 °C, while air ventilation was at 30%. A thermographic camera (FLIR A65, f = 13 mm, lens: 45°; FLIR Systems, Wilsonville, USA) was attached to a metal, height-adjustable universal stand to record from above. A Styrofoam board was placed into the incubator as a standardized and non-reflective background on which the petri dishes with the bees were positioned (for setup see Supplementary Figure S1). Thermographic recording began when the petri dishes with the bees were placed into the incubator. The software FLIR Tools+ v. 6.4.18039.1003 (FLIR Systems, Wilsonville, USA) was used for recording. The emission value was set to 0.97 as this corresponds to the emission of the insect cuticle ^30,31^. We recorded 12 frames/min (every 5 seconds) over a total duration of 22 min and subsequently shock-froze the bees in liquid nitrogen before storing them at -80 °C. The petri dishes were cleaned with 70% EtOH and reused.

### Genotyping and sequencing of the honey bees

Genomic DNA was extracted from two legs using the innuPrep DNA Mini Kit 2.0 (Innuscreen GmbH, Berlin, Germany). We added an RNA digestion step (1 µl RNase A (0.5 u), 5 min incubation at room temperature; Thermo Fisher Scientific, Waltham, USA) on the column-bound DNA to fully remove residues that might have passed the filter.

To detect honey bee mutants, we first performed fluorescent-labeled capillary gel electrophoresis (FCGE) of amplicons covering the sgRNA target site. For this, we used standard Phusion PCR with a reaction volume of 20 µl including a 5’-hexachlorofluorescein-tagged forward primer (for oligonucleotide and amplicon sequences see Supplementary Tables S2, S7). Amplicons were analyzed together with GeneScan™ 500 ROX™ standard (Thermo Fisher Scientific, Waltham, USA) at an ABI 3130XL Genetic Analyzer. Frame shift mutations by insertions or deletions were identified based on length deviation of the amplicons relative to the wild-type fragment length (297 bp) with the Peakscanner cloud software (Thermo Fisher Scientific, Waltham, USA). FCGE analysis indicated n = 57 (41.6%) of the sgRNA-injected honey bees as non-mosaic, biallelic frame shift mutants (mutant type categorization in Supplementary Table S8).

We selected 35 putative *AmOARβ2* knockout mutants for the characterization of the mutations by amplicon deep sequencing with 7 control bees serving as representative wild-type amplicon reference (Supplementary Table S9) and performed a Phusion PCR to create the amplicon spanning the *AmOARβ2* cutting site. Forward primers were composed of the Illumina partial adapter sequence followed by a varying 8 bp-long tag sequence for duplex analysis, and finally the same target-specific hybridization sequence as for the amplicon produced for FCGE. The reverse primer consisted of the sequencing adapter and the target-annealing part (for oligonucleotide and amplicon sequences see Supplementary Tables S2, S7). We used 50 µl reaction volume in the PCR at standard conditions and performed 35 cycles with hybridization at 63 °C for 30 seconds. Amplicons were purified (Monarch Spin PCR & DNA Cleanup Kit; NEB, Ipswich, USA), quantified and merged in equal amounts for NGS analysis in duplexes. Paired-end sequencing was performed on a 2 x 250 Illumina MiSeq device (Illumina, San Diego, USA). Sufficient coverage of the four allele amplicons in a duplex was guaranteed by at least 90,000 total reads. Reads were separated based on their tag sequences and counted using a custom Python script to specify both allele variants of each honey bee. Low-frequency unrelated sequences were removed, while true allele sequences could be reliably determined due to the high read representation.

### Data analysis and statistics

Data analyses were performed with R v. 4.5.0 ^32^ and RStudio v. 2025.09.2+418 ^33^. For data management and visualization, we used tidyverse packages, i.e. dplyr v. 1.1.4 ^34^, readr v. 2.1.5 ^35^, tidyr v. 1.3.1 ^36^, purrr v. 1.0.4 ^37^, stringr v. 1.5.1 ^38^, magrittr v. 2.0.3 ^39^, rstatix v. 0.7.2 ^40^ and forcats v. 1.0.0 ^41^, combined with the plotting packages ggplot2 v. 3.5.2 ^42^, patchwork v. 1.3.0 ^43^, and ggpubr v. 0.6.0 ^44^. Geographic data was visualized using QGIS v. 3.40.13 ^45^. The main map is based on Sentinel-2 cloudless imagery containing modified Copernicus Sentinel data ^46^ and the country boundaries for the overview map were obtained from Natural Earth ^47^.

For quantification of *AmOAR* mRNA expression, we used the package easyqpcr2 v. 0.1.0 ^48^. Data were tested for normal distribution by Shapiro-Wilk normality test. Since data was not distributed normally across all genes and tissues, Mann-Whitney U tests were performed. Raw HPLC data were processed based on the internal standard (IS) and the total brain protein. We created calibration models for the IS and quantified the octopamine. The normalized octopamine concentration in the brains of the bees were determined, accounting for minimal dissection variation by normalization on total protein. The difference in octopamine brain concentration was tested with Mann-Whitney U tests, because data were not normally distributed.

Fisher’s exact tests were used to compare the survival rates of larvae and adult bees between treatments during artificial rearing. After the thermogenesis test, the SEQ files recorded with the thermography camera were first converted to RAW format with the help of the R packages igraph v. 2.2.1 ^49^, imager v. 1.0.5 ^50^, Thermimage v. 4.1.3 ^51^ and fields v. 16.3.1 ^52^. The converted files were loaded into ImageJ v. 1.53k ^53^ with the following parameters: 32-bit-real, 640-pixel width, 512-pixel height, little endian bite order, use virtual stack. Temperatures were measured with the ROI manager for all stack slices (one picture per 5 seconds). An ellipsoid tool was selected and placed over the petri dish covering the entire accessible area for the honey bee tested. With the chosen option min and max gray value, the maximum temperature values in the area were extracted for the observation period. Initially, the bees constituted the coldest area of the assay, as they were transferred directly from ice. Consequently, the first minute of the experiment was excluded from data analysis. Subsequent to this period of torpor, the bees exhibited an increase in body temperature as they underwent thermogenesis. The temperature of their thorax increased to a level that was the highest in the petri dish. The heating phases were analyzed in detail for each bee, with the aim of mirroring their individual heating behavior in a standardized way. The recorded maximum temperature time series values were used to calculate the smoothed temperature using a centered moving average. For each time point, the mean temperature within a 15-measurement window (which corresponds to 75 seconds) was determined. Edge values were computed using the largest available window. For smoothing, the rollapply function in the R package zoo v. 1.8.14 ^54^ was applied. We set three conditions for a temperature increase to be classified as a heating phase: i) a start threshold of > 0.1 °C between two time points, followed by a temperature increase until reaching ii) an end threshold of 0.03 °C where temperature increase reached an asymptote, and iii) an overall temperature change ≥ 1 °C. The comparison of the heating phase characteristics, temperature difference (amplitude) and the heating performance over time during heating phases (slope; see Figure 3 A) was performed by calculation of the weighted means using the general definition

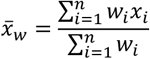

where *x*_*i*_ is the value of the *i*-th heating phase (amplitude or slope, respectively) with the corresponding weight *w*_*i*_ (amplitude). This approach ensured that heating phases with larger amplitude contributed proportionally more to the mean estimate than phases with smaller amplitudes. Weighted mean amplitude and slope were tested with the Mann-Whitney U test for non-normally distributed data. Further characteristics related to thermogenesis were determined: the maximum temperature a tested individual reached during the experiment, the number of heating phases, the weighted mean duration of those heating phases (calculated with the aforementioned formula), the total duration and amplitude of all heating phases, and the area under the curve (AUC, integral of the temperature difference over time during the experiment with the ambient cold stress temperature of 18 °C set as baseline). These characteristics were analyzed for normal distribution with the Shapiro-Wilk normality test and, based on presence of absence of normal distribution, tested with Student’s t-test or Mann-Whitney U test, respectively.

Data were visualized in boxplots displaying the median (horizontal line), the interquartile range (IQR, box), and the individual data points. Statistical significance is indicated by asterisks (*α* = 0.05). Throughout the manuscript, median values are reported with the median absolute deviation (MAD), whereas mean values are specified with the standard error of the mean (SEM).

## Results

### Elevation-dependent octopaminergic signaling in honey bees

Expression of the five octopamine receptor genes differed markedly between honey bees collected at high and low elevations. Tissue-resolved analyses revealed pronounced heterogeneity in these responses: both the direction and magnitude of expression shifts varied across receptor genes and tissues, but one receptor was strikingly different across tissues.

In the brain, honey bees from high elevation exhibited significantly lower expression of both octopamine α receptors and one β receptor (*AmOARβ1*) compared to bees from low elevation (Figure 2 A-C). The other two β receptors did not show differential expression in the brain (Figure 2 D-E, Table 1). In the thorax muscle, the tissue responsible for thermoregulation, mRNA expression of all three octopamine β receptors was significantly increased at high elevation (Figure 2 H-J), while that of the octopamine α receptors did not differ (Figure 2 F-G, Table 1). In the fat body, which has an important function in metabolite storage, only *AmOARβ2* was more highly expressed in bees at higher altitude (Figure 2 N), whereas the remaining receptors did not show expression differences (Figure 2 K-M, O, Table 1). Across all tissues, *AmOARβ2* was the gene showing the most consistent elevation-associated differences in gene expression. Both thermogenesis-related tissues, i.e. thorax muscle and fat body, displayed a significantly higher *AmOARβ2* expression in bees from high elevation. In addition, bees from high elevation contained significantly higher octopamine concentrations in the brain (24.2 ± 6.6 pmol/mg protein) than bees from low elevation (17.0 ± 6.4 pmol/mg protein; Figure 1 B, Table 1).

**Figure 2.**
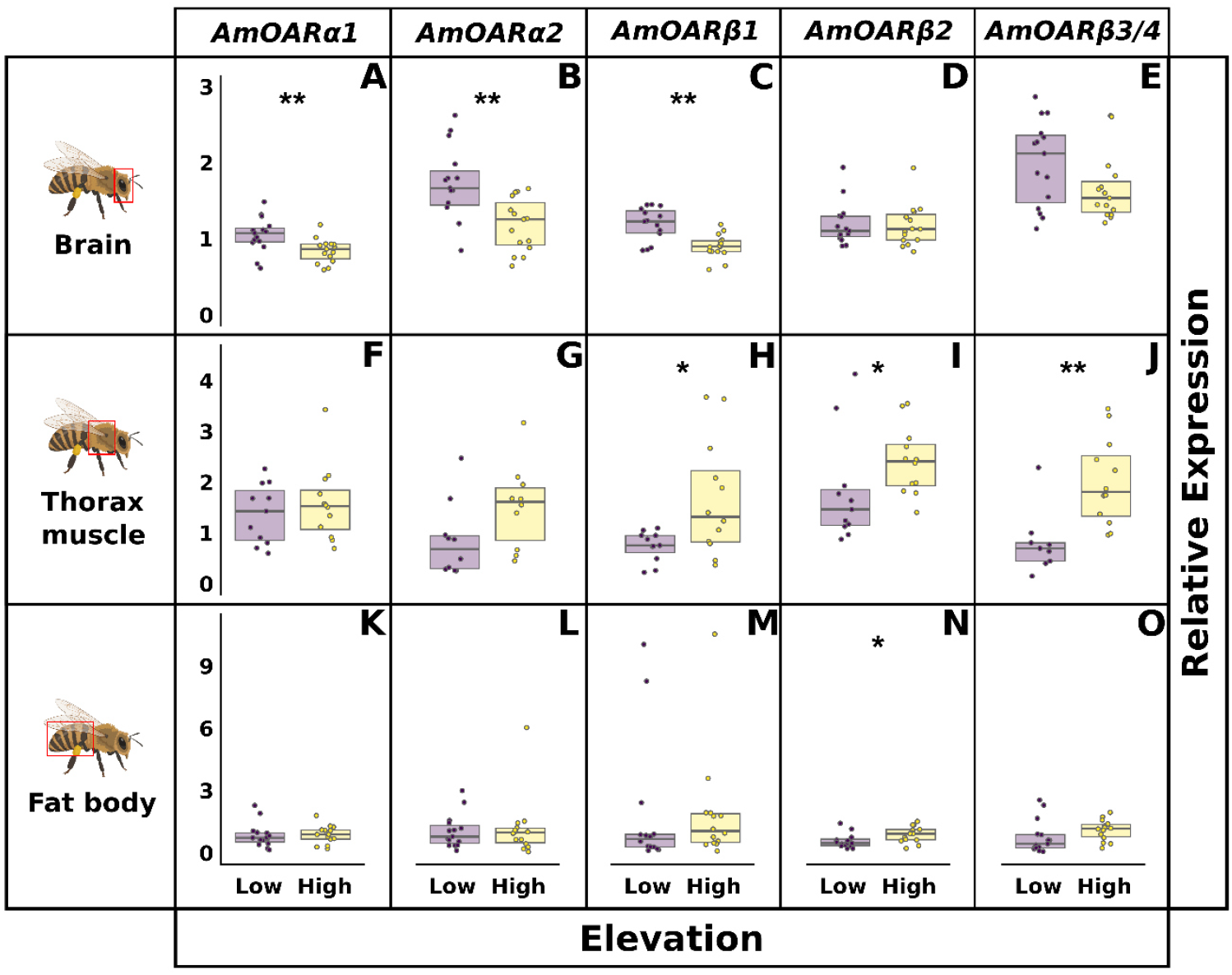
Relative octopamine receptor mRNA expression in honey bees from low elevation (purple) differs markedly from those from high elevation (yellow). Top row: brain (**A-E**). Middle row: thorax muscle (**F-J**). Bottom row: fat body (**L-O**). N = 15 each with some groups showing filtering-dependent lower sample size. Significant differences are indicated by asterisks: * = *P* < 0.05, ** = *P* < 0.01. For detailed statistics see Table 1. Partially created in BioRender, see ^55^.

**Table 1.**
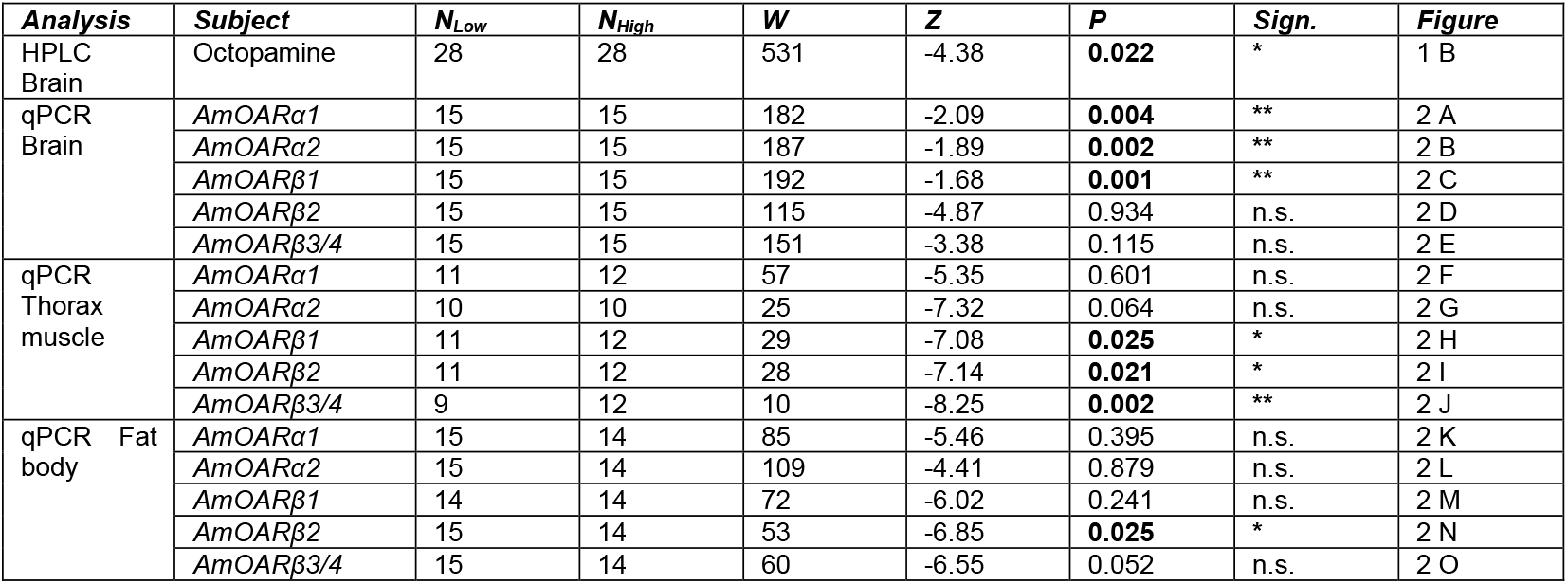
Statistical analysis (Mann-Whitney U test) of the octopamine receptor gene expression relative to reference genes and the octopamine brain concentrations. Significance: * = *P* < 0.05, ** = *P* < 0.01, n.s. = *P* ≥ 0.05 (not significant).

### Loss of *AmOARβ2* altered thermogenesis

Knockout of the octopamine β receptor gene *AmOARβ2* led to a significantly reduced slope of heating phases compared to that of wild type control bees (*OARβ2*^*-/-*^ vs. *OARβ2*^*wt/wt*^; Table 2, Figure 3 B), suggesting a reduced heat production under cold stress. Given the similar average length of heating phases, this resulted in a reduced heating amplitude in *OARβ2*^*-/-*^ mutants (Table 2, Figure 3 C). They demonstrated an approximately 2 °C lower average amplitude than wild-type honey bees when heating. These findings demonstrate an altered dynamics in thermogenic behavior in mutants lacking *AmOARβ2*.

**Table 2.** Statistical analysis of the heating phase characteristics in the cold stress assay. Comparison of mutants (*OARβ2*^*-/-*^) and wild-type bees (*OARβ2*^*wt/wt*^). Median (M) followed by median absolute deviation (MAD), mean followed by standard error of the mean (SEM) Significance: * = *P* < 0.05, ** = *P* < 0.01, n.s. = *P* ≥ 0.05 (not significant).

| Subject | $M \pm$<br>$MAD_{OAR\beta 2^{-/-}}$ | $M \pm$<br>$MAD_{OAR\beta 2^{wt/wt}}$ | $N_{OAR\beta 2^{-/-}}$ | $N_{OAR\beta 2^{wt/wt}}$ | Test | W | Z | P | Sign. |
| --- | --- | --- | --- | --- | --- | --- | --- | --- | --- |
| Weighted mean slope | $0.025 \pm 0.009$ °C/s | $0.038 \pm 0.015$ °C/s | 35 | 38 | Mann-Whitney U test | 374 | -10.18 | **0.001** | **\*\*** |
| Weighted mean amplitude | $4.4 \pm 3.2$ °C | $6.2 \pm 3.0$ °C | | | | 486 | -8.94 | **0.048** | **\*** |
| Weighted mean duration | $171 \pm 80$ s | $163 \pm 50$ s | | | | 702 | -6.55 | 0.687 | n.s. |
| Number of heating phases | $2 \pm 1.5$ | $2 \pm 1.5$ | | | | 677 | -6.83 | 0.897 | n.s. |
| Maximum temperature | $29.3 \pm 7.7$ °C | $31.1 \pm 6.9$ °C | | | | 607 | -7.60 | 0.525 | n.s. |
| AUC | $7365 \pm 5053$ °C·s | $8515 \pm 4143$ °C·s | | | | 638 | -7.26 | 0.766 | n.s. |
| Subject | $Mean \pm$<br>$SEM_{OAR\beta 2^{-/-}}$ | $Mean \pm$<br>$SEM_{OAR\beta 2^{wt/wt}}$ | $N_{OAR\beta 2^{-/-}}$ | $N_{OAR\beta 2^{wt/wt}}$ | Test | t | Df | P | Sign. |
| Total duration | $308 \pm 26$ s | $280 \pm 19$ s | 35 | 38 | Student's t-test | 0.86 | 71 | 0.395 | n.s. |
| Total amplitude | $8.4 \pm 0.9$ °C | $10.3 \pm 1.0$ °C | | | | -1.38 | 71 | 0.172 | n.s. |

**Figure 3.**
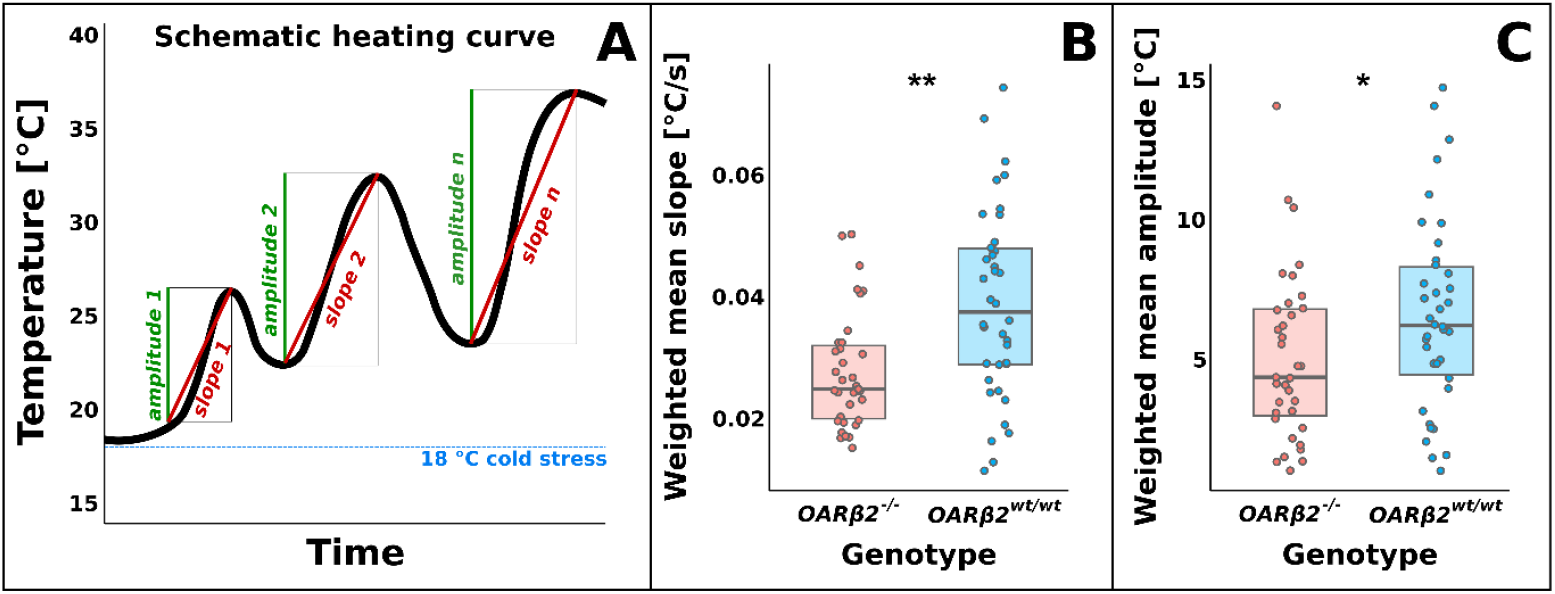
Thermogenesis of mutants lacking the octopamine β2 receptor and wild-type control bees during cold stress. **A**. Schematic example of heating curve during the experiment. The honey bees showed individual temperature profiles with *n* heating phases from which we extracted slope and amplitude. **B**. Weighted mean slope calculated from the slopes of all heating phases for *OARβ2*^*-/-*^ mutants (red) and wild-type bees (*OARβ2*^*wt/w*t^, blue). **C**. Weighted mean amplitude calculated from the amplitudes of all heating phases for *OARβ2*^*-/-*^ mutants (red) and wild type bees (*OARβ2*^*wt/wt*^, blue). N (*OARβ2*^*-/-*^) = 35, N (*OARβ2*^*wt/wt*^) = 38, significant differences are indicated by asterisks: * = *P* < 0.05, ** = *P* < 0.01. For detailed statistics see Table 2.

Interestingly, loss of *AmOARβ2* did not block thermogenesis completely, suggesting a modulatory function of this octopamine receptor rather than an eliciting function. Overall heating performance was generally comparable in wild-type bees and mutants over the entire duration of cold stress exposure (area under the curve; Table 2). The number, total duration and amplitude of all heating phases did not differ between the two treatments (Table 2). Under 18 °C cold stress, both mutants and wild types actively increased their thorax temperature substantially and reached a maximum of around 30 °C. Mutants had a slightly lower maximum thorax temperature with 29.3 ± 7.7 °C compared to wild types (31.1 ± 6.9 °C; Table 2).

## Discussion

### Octopaminergic signaling in thermogenic tissue depends on elevation

A central finding of our study is the elevation-dependent regulation of octopamine signaling in thermogenic tissues. In the thoracic flight muscle and abdominal fat body, the mRNA expression of all three β-adrenergic octopamine receptor genes was significantly elevated in bees from high elevation compared to those from low elevation. This provides the first direct evidence for the hypothesis that octopamine β receptors are involved in adaptation to elevation, which was suggested by Wallberg et al. ^21^ based on chromosomal inversions in bees from high and low elevations on Mt Kenya. The assumption that octopamine β receptors mediate thermogenesis in honey bees was recently suggested by Kaya-Zeeb et al. ^10,11^. Our behavioral tests on thermogenesis performance of honey bees lacking the octopamine β2 receptor under cold stress directly confirm this hypothesis.

All octopamine β receptors are located in the divergent inversion polymorphism region on chromosome 7, suggesting that they are under selection for octopamine signaling in high elevation. Structural inversions have been shown to facilitate local adaptation in various species, thereby sustaining divergent ecotypes and adaptive phenotypes ^56–58^. For honey bees, evidence for divergent highland haplotypes in East Africa carrying an inversion involving the octopamine β receptors is not based solely on one study from Mt. Kenya ^21^. Further mountains in East Africa, like Mt. Elgon in Kenya ^59,60^ and Rwenzori mountains investigated by our group in Uganda ^23^, accommodate high-elevation adapted honey bees which differ from those at lower elevations. Regardless of slight variations in the inversion across mountain systems, the octopamine β receptor-harboring genomic regions are apparent for highland adaptation in Ugandan honey bees, too ^23^. Our findings thus confirm at the transcriptomic level that honey bees living in higher elevation habitats adapt their amine signaling system by increased octopamine β receptor expression, while the impact of octopamine α receptors on adaptation to elevation seems less clear.

Further functions of octopamine β receptors have not been described in honey bees yet. Their characterization in the honey bee was only reported about a decade ago, based on brain tissues ^14^. Since then, few studies have investigated specific octopamine β receptors in tissues beyond the brain. Besides an elevation-associated downregulation of *AmOARβ1* in the head ^23^, expression differences of *AmOARβ3/4* in the mushroom bodies of honey bee brains were associated with age rather than social role ^61^. Pharmacological manipulation of octopamine levels including elevation, depletion and non-specific receptor antagonism has produced diverse effects ^5,8,9,14,19,20^. Yet the contributions of individual receptors in specific traits and behaviors of honey bees deserve further investigation. The upregulation of *AmOARβ2* in the fat body might reflect the modulation of metabolism-associated processes, because octopamine can affect energy mobilization and arousal ^13^. In fruit flies, mutants lacking *Octβ1R* or *Octβ2R* displayed obesity phenotypes similar to flies lacking the transmitter octopamine ^62^. *Octβ1R* was proven to be directly involved in fat-body catabolism ^63^, whereas the function of *Octβ2R* in fat body lipid mobilization is unclear.

A notable finding is the differential expression of *AmOARβ1* in the brain of bees collected at high and low elevations in our current study. This finding directly corroborates the results of our earlier experiments conducted on honey bees at the Rwenzori mountain slopes in Western Uganda. There, we also found *AmOARβ1* to be downregulated in the head tissues of bees from high elevation ^23^. The low elevation in our earlier study was nearly identical to our current low elevation, and the high elevation collection site was with > 2,000 m ASL similar to our current high elevation sites. Although the exact function of *AmOARβ1*-mediated signaling in the brain and its role in adaptation to elevation has yet to be studied, we speculate that this receptor might have a regulatory function in sensory information processing or in responsiveness to external stimuli relevant for honey bees at high elevation.

In contrast to the prominent upregulation of the octopamine β receptors in thorax muscle and fat body at high elevation, we report a decreased expression of the two α-adrenergic receptor genes *AmOARα1* and *AmOARα2* in the brain. The underlying reasons for this divergent expression pattern in comparison to the β receptors remains speculative, as the respective proteins may be involved in a multitude of processes. AmOARα1 is widely present throughout the brain and has been linked to the processing of sensory information, learning and memory ^64,65^. To date, no correlation to thermoregulation or metabolism has been identified. *AmOARα2* has been characterized pharmacologically only recently, and there is no information regarding its function in honey bees available ^15^. Both octopamine α receptors are not located within the genomic chromosomal inversion region. They differ in modulation action and are linked to different signaling cascades (AmOARα1 evokes Ca^2+^ release upon activation, AmOARα2 decreases cAMP by adenylyl cyclase inhibition). This indicates a complex and fine-tuned modulatory system that remains to be fully understood.

In addition to transcriptomic differences between bees from high and low elevations, octopamine concentrations were higher in the brains of honey bees from high elevation. The honey bee brain is the center of sensory input and information processing and therefore highly sensitive to biogenic amine signaling ^9,13,26,66^. Interestingly, the octopamine concentration in the brain of worker bees was earlier shown to decrease under cold stress ^67^. Lower ambient temperatures at higher elevation imply enhanced cold stress for honey bees. Honey bee foragers need to raise their body temperature before flight ^7,30,68^. If we consider high elevation as a proxy for cold stress, we find the opposite pattern than hypothesized by Chen et al. ^67^. High-elevation honey bees might have gone through an altitudinal adaptation process such as was reported for the octopamine receptor genes in East African honey bees ^21^. Adaptation to cold stress at high elevation might be constrained by hitherto unknown factors which are tightly linked to elevation. Given the high pleiotropy of octopamine modulation in the bee ^8,13,20^, elevated octopamine levels may arise from further interacting, octopamine-dependent processes. This could possibly involve landscape complexity, associated information processing and memory formation to perform adequate foraging behavior ^19,66,69^. Furthermore, higher elevation is associated with lower partial pressure of oxygen. Octopamine has been shown to function as a compensatory neuromodulator that buffers hypoxia-induced impairments in neuronal signaling in locusts ^70^, suggesting a similar role in honey bees.

Environmental conditions are strikingly different at mountain clines. The “stress hormone” octopamine ^20^ and its receptors may therefore be of major significance for adaptation. Temperature is one of the most notable impacts on insect physiology at high elevations ^71^. With every 1,000 m of altitudinal increase along a mountain slope there is an approximate temperature reduction of 5.2 °C – 6.5 °C ^72^. Thus, we hypothesize that the observed octopamine-related differences are connected to thermal adaptation. G-protein-coupled receptors like the octopamine receptors are conserved throughout insects with closely related gene orthologues ^12^. However, evidence for an octopamine receptor modulating thermogenesis from similarly endothermic taxa like bumble bees or moths is lacking.

Our experiments with *AmOARβ2* knockouts confirm the importance of this receptor gene for thermoregulation. Although total heat production was comparable, mutants displayed a reduced rate of heating performance during the heating events. Consequently, despite heating phases of similar duration, they reached a lower average temperature amplitude. These findings imply that octopamine may have a modulatory function on thermoregulation rather than initiating heat production directly by activating wing muscles. Intriguingly, we can demonstrate that the loss of *AmOARβ2* did not impair thermogenesis completely. This presents a new perspective on intertwined modulation, challenging the previously proposed hypothesis that at least one isoform of *AmOARβ2* is essential for cold stress response by thermogenesis in honey bees ^10^. Thermogenesis in mutant honey bees lacking functional AmOARβ2 emphasizes that additional genes must be involved, potentially other octopamine β receptors acting through the same signaling cascade. Given the adaptive importance of thermogenesis for both individual honey bees and the colony, a compensatory mechanism regulating this central behavior seems likely. Similarly, honey bees lacking a highly specific fructose receptor still responded to this sugar with proboscis extension, although the proportion of responders was very low, indicating a comparable backup system ^73^.

Connecting all of our results, we suggest that the increased octopamine signaling reflects a well-adapted response to high-elevation thermal conditions. It has been shown that octopamine modulates permanent adaptation in reaction to chronic stimuli ^63,74^. If we transfer this to chronic cold stress, we hypothesize that the increased receptor mRNA expression indicates an adaptive trait for high-elevation adaptation, supporting the steady-state hypothesis proposed for the octopaminergic system ^10^. A constant synthesis and availability of high concentrations of octopamine, and numerous octopamine receptors could facilitate adaptive responses to thermal stress at high elevations.

### Conclusion

Our findings strongly indicate that octopaminergic signaling contributes to thermal performance in cold environments. The differences in brain octopamine levels and tissue-dependent octopamine receptor gene expression suggest a coordinated modulation of neural and peripheral components of the thermogenic system. Together with the altered heating behavior in *AmOARβ2* knockouts, this supports a key role for this signaling pathway in high-elevation adaptation.

The world is heating up, and global change scenarios indicate a future with increasing temperature and more extreme events ^75^. If species fail to adapt to changing conditions, this could result in forced range shifts, reduced distribution and population sizes, and ultimately an increased risk of extinction. Honey bees with their remarkable thermoregulatory strategies are able to buffer thermal stress from the environment to a larger extent than most other insects. However, considering the environmental and economic importance of *Apis mellifera* in Kenya and on a global scale ^76,77^, further research is crucial to fully understand what contributes to successful local adaptation in a changing world. Mountains constitute ideal study sites because of the gradual variation of environmental conditions along the altitudinal cline. The octopaminergic system with its pleiotropic impact on physiological and behavioral processes is a key candidate for future research trying to understand elevation-driven, local adaptation in honey bees.

## Resource availability

### Lead contact

Further information and requests for resources should be directed to, and will be fulfilled by, the lead contacts, Florian Loidolt and Ricarda Scheiner.

### Materials availability

This study did not generate new, unique materials.

### Data and code availability

- All datasets and code are publicly available via https://doi.org/10.5281/zenodo.19626461 on Zenodo.
- Any additional information required to reanalyze the data reported in this paper is available from the lead contact upon request.

## Acknowledgments

We thank Kennedy Talam for his incredible help during Kenyan fieldwork, communication with local beekeepers and honey bee sampling. We thank the Kenyan collaborating beekeepers and their families, kindly supporting our research: John Nyagah Munya, Mercy Muthanje, Moses Rugendo, Faith Muthoni and Amos Mureithi Njue from Rukuriri area, and Benjamin Kagundu from Kiritiri. We thank the University of Embu as collaborating research facility, especially Prof. Romano Mwirichia. We thank the whole University of Würzburg CRISPR team, involved beekeepers and lab technician personnel. We thank Markus Ankenbrand and the HackyHour team for help in data analysis. This research was supported by grants of the Deutsche Forschungsgemeinschaft (DFG) to R.S. and M.H. (SCHE 1573/14-1; HA 5499/11-1) and by a grant of the Volkswagenstiftung to R.S.

## Author contributions

R.S. and M.H. equally managed lead conceptualization, funding acquisition, administration and supervision in the shared project; F.L. and M.M. performed the Kenyan fieldwork and honey bee sampling in great collaboration with M.O.; F.L. did the CRISPR/Cas9 mutant generation and thermogenesis experiments and performed the laboratory analyses in coordination with M.T.; F.L. did the lead data analysis for with input from M.T. and R.S.; F.L. had the lead in visualization and writing (original draft, review and editing), with all authors supporting during the review and editing process.

## Declaration of interest

The authors declare no competing interests.

## Supplemental information

- Supplementary Tables S1-S9
- Supplementary Figure S1

## Notes

https://doi.org/10.5281/zenodo.19626461

### Competing Interest Statement

The authors have declared no competing interest.

