## Supplementary Tables and Figures for "Octopaminergic signaling contributes to thermal adaptation to elevation in African honey bees (*Apis mellifera*)"

### Supplementary Data

#### Supplementary Figure S1

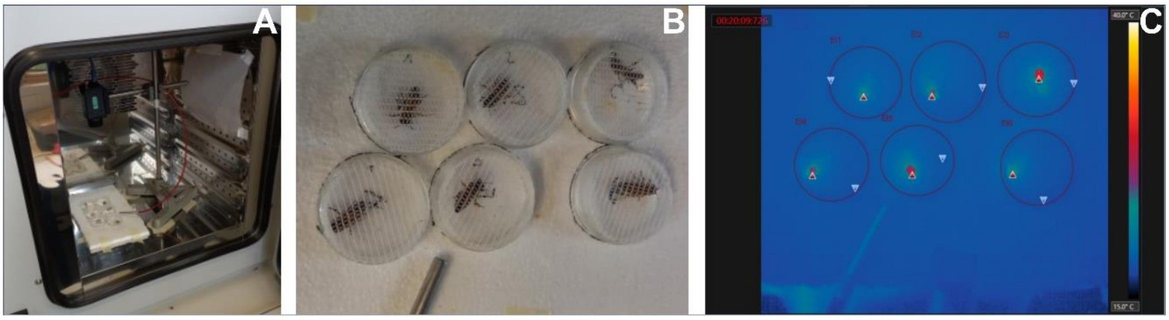


**Figure S1. Behavioral cold stress assay to investigate thermogenic performance.** Worker bees (fed with 50% sucrose solution and immobilized on ice) were placed in petri dishes with 3 cm diameter (6 bees tested per trial) and recorded with a thermographic camera under 18 °C cold stress. **A.** Experimental setup in the incubator device with a thermographic camera mounted on a metal stand to record from above. **B.** Tested bees in petri dishes. **C.** Visualization of the heating behavior of the honey bees. In each petri dish (circles), the honey bees' thoraces were the area of maximum temperature (red triangles) indicating thermogenesis, while the coldest spots (blue triangles) were always located apart from the bees.

#### Supplementary Tables S1 – S9

##### Table S1. Sample site information at Mt. Kenya high and low elevations.

| ***Colony ID*** | ***High/Low*** | ***Elevation [m ASL]*** | ***Latitude (DD)*** | ***Longitude (DD)*** | ***Samples HPLC*** | ***Samples qPCR*** |
|---|---|---|---|---|---|---|
| HL1 | High | 1910 | -0.3555 | 37.49582 | 10 | 5 |
| HL2 | High | 1910 | -0.3555 | 37.49582 | 10 | 5 |
| HL3 | High | 1846 | -0.36768 | 37.50343 | 10 | 5 |
| LL1 | Low | 1161 | -0.68436 | 37.6366 | 10 | 5 |
| LL2 | Low | 1162 | -0.68476 | 37.63645 | 10 | 5 |
| LL3 | Low | 1163 | -0.68495 | 37.63628 | 10 | 5 |

**Table S2. Oligonucleotides used in the study.** qPCR SYBR = Quantitative real-time PCR with SYBR Green, for gene expression analysis. FCGE = Fluorescent capillary gel electrophoresis, for determination of the amplicon length. Amplicon SEQ = Amplicon sequencing, for CRISPR/Cas9-induced mutation specification; Amplicon SEQ primers contained the Illumina partial adapter sequences (italic) and the forward primers had specific tags (underlined) for duplex analysis.

| ***Analysis*** | ***Gene*** | ***Gene ID*** | ***Direction*** | ***Sequence (5'-3')*** |
|---|---|---|---|---|
| qPCR SYBR | *AmOARα1* | 406068 | forward | GCAGGAGGAACAGCTGCGAG |
| | | | reverse | GCCGCCTTCGTCTCCATTCG |
| qPCR SYBR | *AmOARα2* | 726331 | forward | GCGAGCATCATGAACTTGTG |
| | | | reverse | CGTAGCCTATGTCCTCTGAAAG |
| qPCR SYBR | *AmOARβ1* | 413698 | forward | GGAGTAAAGTAGCAGCCGCTC |
| | | | reverse | GTGATCTGTGGCTCCTCTGGT |
| qPCR SYBR | *AmOARβ2* | 412896 | forward | CTCGAGCGAGGAGAAGTTGT |
| | | | reverse | CCAACGCTAAAGAGACCACG |
| qPCR SYBR | *AmOARβ3/4* | 412994 | forward | CGAGGACGCTCGGAATAATA |
| | | | reverse | GAAGTCGCGGTTGAAGTACG |
| qPCR SYBR | *AmRpL10* | 409589 | forward | CGATAAGAAACGTAAGTCAATATGGGGC |
| | | | reverse | CGTATCTTTGGATCAGGCACACC |
| qPCR SYBR | *AmRpL19* | 724186 | forward | GGGACTTCTAGGCTCCATCATGAG |
| | | | reverse | GCTTTGACGTGAGTTTGTATTTGCAA |
| qPCR SYBR | *AmRpL32* | 406099 | forward | AGTAAATTAAAGAGAAACTGGCGTAA |
| | | | reverse | TAAAACTTCCAGTTCCTTGACATTAT |
| FCGE | *AmOARβ2* | 412896 | forward | HEX-CCTCGGTGGACGTGACG |
| | | | reverse | CGGTCAATTGCACGCTC |
| Amplicon SEQ | *AmOARβ2* | 412896 | forward tag A | *ACACTCTTTCCCTACACGACGCTCTTCCGATCT*CTGTGATGCCTCGGTGGACGTGACG |
| | | | forward tag B | *ACACTCTTTCCCTACACGACGCTCTTCCGATCT*GCGCAATACCTCGGTGGACGTGACG |
| | | | reverse | *GACTGGAGTTCAGACGTGTGCTCTTCCGATCT*CGGTCAATTGCACGCTC |

**Table S3. sgRNA candidate information and selection for AmOARβ2 knockout based on test injections.** CDS = coding DNA sequence, PAM = protospacer adjacent motif, 24 h mortality comprises the injection capabilities and the egg's tolerance for the needle penetration and wound repair, larval hatching rate [%] = (hatching larvae / (injected eggs - 24 h mortality)) x 100, mutation frequency considers only full knockout larvae with two frame shift mutation alleles based on fluorescent capillary gel electrophoresis for determination of the amplicon length, mutant rate [%] = larval hatching rate x mutation frequency, colored sgRNA candidate and concentration were selected for the experimental injections based on the highest mutant rate.

| ***General information*** | ***sgRNA-candidate 1*** | | | ***sgRNA-candidate 2*** | | | ***sgRNA-candidate 3*** | | |
|---|---|---|---|---|---|---|---|---|---|
| guide sequence 5'-3' | GGTCGAGATGCCGTTCAACA | | | GAACGCGAGCTACTCGAGCG | | | GAACATGGCGACCAACATGT | | |
| CDS location & strand | 68 - | | | 122 + | | | 290 - | | |
| PAM | GGG | | | AGG | | | CGG | | |
| on-target score | 65.3 | | | 67.2 | | | 69.1 | | |
| off-target score | 94.9 | | | 96.1 | | | 98.2 | | |
| ***Test injection results*** | ***Concentration 1*** | ***Concentration 2*** | ***Concentration 3*** | ***Concentration 1*** | ***Concentration 2*** | ***Concentration 3*** | ***Concentration 1*** | ***Concentration 2*** | ***Concentration 3*** |
| sgRNA concentration [ng/µl] | 23 | 46 | 92 | 23 | 46 | 92 | 23 | 46 | 92 |
| Cas9 concentration [µM] | 3.13 | 3.13 | 3.13 | 3.13 | 3.13 | 3.13 | 3.13 | 3.13 | 3.13 |
| injected eggs | 295 | 219 | 418 | 378 | 182 | 311 | 215 | 217 | 175 |
| 24 h mortality | 170 | 73 | 234 | 275 | 43 | 205 | 74 | 99 | 80 |
| larval hatching rate [%] | 11.2 | 11.6 | 8.7 | 21.4 | 14.4 | 15.1 | 24.1 | 19.5 | 17.9 |
| genotyped larvae | 14 | 13 | 6 | 12 | 5 | 14 | 15 | 14 | 14 |
| mutation frequency [%] | 15 | 23 | 0 | 17 | 40 | 57 | 7 | 7 | 7 |
| mutant rate [%] | 2 | 3 | 0 | 4 | 6 | 9 | 2 | 1 | 1 |

**Table S4. Honey bee eggs used for the knockout experiment, 24 h survival and larval hatching for the three replicate injection days.** Larval hatching rate refers to successful eclosion from eggs which were alive after 24 hours.

| | ***# eggs*** | | | ***24 h survival [# (%)]*** | | | ***Larval hatching [# (%)]*** | | |
|---|---|---|---|---|---|---|---|---|---|
| ***Treatment*** | ***Day 1*** | ***Day 2*** | ***Day 3*** | ***Day 1*** | ***Day 2*** | ***Day 3*** | ***Day 1*** | ***Day 2*** | ***Day 3*** |
| injected | 683 | 838 | 574 | 487 (71.3) | 582 (69.5) | 411 (71.6) | 193 (39.6) | 149 (25.6) | 123 (29.9) |
| control | 25 | 98 | 73 | 24 (96.0) | 94 (95.9) | 72 (98.6) | 24 (100) | 93 (98.9) | 70 (97.2) |

**Table S5. Survival of the worker bees reared in the laboratory from larval stage until adult eclosion.**

| *Treatment* | *# of hatched larvae used for rearing* | *# of eclosed adult honey bees* | *survival to adult stage [%]* | *Fisher's exact test, P-value, df = 1* |
|---|---|---|---|---|
| injected | 370 | 211 | 57.0 | 0.85 |
| control | 159 | 89 | 56.0 | |

**Table S6. Survival of the eclosed worker bees until reaching the experimental age of 14 days.** * Considered are fewer eclosed adult bees than listed in Supplementary Table S6 because bees that had emerged much earlier (several days prior than the calculated eclosion period) were excluded.

| *Treatment* | *# of eclosed adults ** | *# of 14 d old honey bees* | *survival until the test age of 14 days [%]* | *Fisher's exact test, P-value, df = 1* |
|---|---|---|---|---|
| injected | 198 | 176 | 88.9 | 0.53 |
| control | 88 | 81 | 92.0 | |

**Table S7. *AmOARβ2* amplicon sequences.** The amplicon for fluorescent capillary gel electrophoresis (FCGE) was designed with a length of 297 bp and includes the CRISPR/Cas9 manipulation site. The amplicon for sequencing (SEQ) was designed with a length of 370 bp, including the same 297 bp fragment, additional Illumina adapters, and tags for duplex analysis. Color code: purple = primer hybridization motifs, blue = sgRNA target sequence with the double strand break site indicated in red, green = Illumina partial sequencing adapters, orange = 8 bp tag sequence (see Supplementary Table S2).

<table>
<tr><th>Analysis</th><th>Sequence (5'-3')</th></tr>
<tr><td>FCGE</td><td>HEX-<br>CCTCGGTGGACGTGACGACCCTGTTGAACGGCATCTCGACCGAGGACGGCCAGCTGGGAACGAACGCG<br>AGCTACTCGA||GCGAGGAGAAGTTGTCGGTGCCTGTGACGATCGTGAAAGGTTGCGTGTTGGGCTCCATC<br>ATAGTGACCGCGGTGTTCGGCAATCTGTTGGTGATGGTGTCGGTGATGCGGCACAGGAAGCTGCGGATCA<br>TCACCAATTATTTCGTGGTCTCTTTAGCGTTGGCCGACATGTTGGTCGCCATGTTCGCCATGACGTTCAACG<br>CGAGCGTGCAATTGACCG</td></tr>
<tr><td>SEQ</td><td>ACACTCTTTCCCTACACGACGCTCTTCCGATCTNNNNNNNNCCTCGGTGGACGTGACGACCCTGTTGAAC<br>GGCATCTCGACCGAGGACGGCCAGCTGGGAACGAACGCGAGCTACTCGA||GCGAGGAGAAGTTGTCGGT<br>GCCTGTGACGATCGTGAAAGGTTGCGTGTTGGGCTCCATCATAGTGACCGCGGTGTTCGGCAATCTGTTG<br>GTGATGGTGTCGGTGATGCGGCACAGGAAGCTGCGGATCATCACCAATTATTTCGTGGTCTCTTTAGCGTT<br>GGCCGACATGTTGGTCGCCATGTTCGCCATGACGTTCAACGCGAGCGTGCAATTGACCGAGATCGGAAGA<br>GCACACGTCTGAACTCCAGTC</td></tr>
</table>

**Table S8. The rate of generating biallelic *AmOARβ2*$^{-/-}$ knockout mutant honey bees.** Mosaic contains bees across all mutation types where more than two different amplicon lengths were detected in fluorescent capillary gel electrophoresis. Other genotype relates to honey bees with mono or biallelic in-frame (if) mutation identified, i.e. *AmOARβ2*$^{wt/if}$, *AmOARβ2*$^{if/if}$ and *AmOARβ2*$^{-/if}$.

| *Genotype* | *adult worker bees [# (%)]* |
|---|---|
| *AmOARβ2*$^{-/-}$ | 57 (41.6) |
| *AmOARβ2*$^{wt/-}$ | 4 (2.9) |
| *AmOARβ2*$^{wt/wt}$ | 24 (17.5) |
| mosaic | 29 (21.2) |
| other | 23 (16.8) |
| **Total** | **137** |

**Table S9. *AmOARβ2* amplicon sequencing mutation confirmation.** The sequenced amplicon located in CDS exon 1 (N-terminal coding exon shared across all transcript variants) starts at the second base of an amino acid (aa) coding triplet, hence we had to consider the triplet structure from the first actual nucleotide triplet on for the detection of frame shift mutations. Therefore, the two starting nucleotides CC (in brackets) were not considered. Early stop codons (red, underlined) occurred due to frame shift mutation. Some mutants did not show early stop codons, but a deviating nucleotide sequence with a frame shift (underlined), resulting in a complete nonsense aa sequence.

Amplicon reference sequence

>ref_seq_AmOAb2R
[CC]TCGGTGGACGTGACGACCCTGTTGAACGGCATCTCGACCGAGGACGGCCAGCTGGGAACGAACGCGAGCTACTCGA
GCGAGGAGAAGTTGTCGGTGCCTGTGACGATCGTGAAAGGTTGCGTGTTGGGCTCCATCATAGTGACCGCGGTGTTCGGC
AATCTGTTGGTGATGGTGTCGGTGATGCGGCACAGGAAGCTGCGGATCATCACCAATTATTTCGTGGTCTCTTTAGCGTTGG
CCGACATGTTGGTCGCCATGTTCGCCATGACGTTCAACGCGAGCGTGCAATTGACCG

Representative wild type

Note: For the wild type, we encountered two amplicon variants during sequencing, one showed a single base change (blue) but without affecting aa composition.

>a44_allele_A, aa 80
[CC]TCGGTGGACGTGACGACCCTGTTGAACGGCATCTCGACCGAGGACGGCCAGCTGGGAACGAACGCGAGCTACTCGA
GCGAGGAGAAGTTGTCGGTGCCTGTGACGATCGTGAAAGGTTGCGTGTTGGGCTCCATCATAGTGACCGCGGTGTTCGGC
AATCTGTTGGTGATGGTGTCGGTGATGCGGCACAGGAAGCTGCGGATCATCACCAATTATTTCGTGGTCTCTCTAGCGTTG
GCCG

>a44_allele_B, aa 80
[CC]TCGGTGGACGTGACGACCCTGTTGAACGGCATCTCGACCGAGGACGGCCAGCTGGGAACGAACGCGAGCTACTCGA
GCGAGGAGAAGTTGTCGGTGCCTGTGACGATCGTGAAAGGTTGCGTGTTGGGCTCCATCATAGTGACCGCGGTGTTCGGC
AATCTGTTGGTGATGGTGTCGGTGATGCGGCACAGGAAGCTGCGGATCATCACCAATTATTTCGTGGTCTCTTTAGCGTTG
GCCG

Mutants

Note: Some honey bee mutants showed the same mutation in both alleles, while other had two different amplicon sequences for each allele (_allele_A & _allele_B).

>a151_allele_A, aa 26
[CC]TCGGTGGACGTGACGACCCTGTTGAACGGCATCTCGACCGAGGACGGCCAGCTGGGAACGAACGCGAGCTACTCGA
AG**TAG**CGAGGAGAAGTTGTCGGTGCCTGTGACGATCGTGAAAGGTTGCGTGTTGGGCTCCATCATAGTGACCGCGGTGTT
CGGCAATCTGTTGGTGATGGTGTCGGTGATGCGGCACAGGAAGCTGCGGATCATCACCAATTATTTCGTGGTCTCTTTAGC
GTTG

>a151_allele_B, aa 32
[CC]TCGGTGGACGTGACGACCCTGTTGAACGGCATCTCGACCGAGGACGGCCAGCTGGGAACGAACGCGAGCTACTCGA
GGAGAAGTTGTCGGTGCCTG**TGA**CGATCGTGAAAGGTTGCGTGTTGGGCTCCATCATAGTGACCGCGGTGTTCGGCAATC
TGTTGGTGATGGTGTCGGTGATGCGGCACAGGAAGCTGCGGATCATCACCAATTATTTCGTGGTCTCTTTAGCGTTGGCCG
ACAT

>a85_allele_A, aa 32
[CC]TCGGTGGACGTGACGACCCTGTTGAACGGCATCTCGACCGAGGACGGCCAGCTGGGAACGAACGCGAGCTACTCGA
GGAGAAGTTGTCGGTGCCTG**TGA**CGATCGTGAAAGGTTGCGTGTTGGGCTCCATCATAGTGACCGCGGTGTTCGGCAATC
TGTTGGTGATGGTGTCGGTGATGCGGCACAGGAAGCTGCGGATCATCACCAATTATTTCGTGGTCTCTCTAGCGTTGGCCG
ACAT

>a85_allele_B, aa 32
[CC]TCGGTGGACGTGACGACCCTGTTGAACGGCATCTCGACCGAGGACGGCCAGCTGGGAACGAACGCGAGCTACTCGA
GGAGAAGTTGTCGGTGCCTG**TGA**CGATCGTGAAAGGTTGCGTGTTGGGCTCCATCATAGTGACCGCGGTGTTCGGCAATC
TGTTGGTGATGGTGTCGGTGATGCGGCACAGGAAGCTGCGGATCATCACCAATTATTTCGTGGTCTCTT**TAG**CGTTGGCCG
ACAT

>b130_allele_A, aa 29
[CC]TCGGTGGACGTGACGACCCTGTTGAACGGCATCTCGACCGAGGACGGCCAGCTGGGAACGAACGCGAGGAGAAGTT

GTCGGTGCCTG**TGA**CGATCGTGAAAGGTTGCGTGTTGGGCTCCATCATAGTGACCGCGGTGTTCGGCAATCTGTTGGTGA
TGGTGTCGGTGATGCGGCACAGGAAGCTGCGGATCATCACCAATTATTTCGTGGTCTCTTTAGCGTTGGCCGACATGTTGG
TCGC

>b130_allele_B, aa 32
[CC]TCGGTGGACGTGACGACCCTGTTGAACGGCATCTCGACCGAGGACGGCCAGCTGGGAACGAACGCGAGCTACTCGA
GGAGAAGTTGTCGGTGCCTG**TGA**CGATCGTGAAAGGTTGCGTGTTGGGCTCCATCATAGTGACCGCGGTGTTCGGCAATC
TGTTGGTGATGGTGTCGGTGATGCGGCACAGGAAGCTGCGGATCATCACCAATTATTTCGTGGTCTCTCTAGCGTTGGCCG
ACAT

>a160_allele_A, aa 32
[CC]TCGGTGGACGTGACGACCCTGTTGAACGGCATCTCGACCGAGGACGGTCAGCTGGGAACGAACGCGAGCTACTCGA
GGAGAAGTTGTCGGTGCCTG**TGA**CGATCGTGAAAGGTTGCGTGTTGGGCTCCATCATAGTGACCGCGGTGTTCGGCAATC
TGTTGGTGATGGTGTCGGTGATGCGGCACAGGAAGCTGCGGATCATCACCAATTATTTCGTGGTCTCTTTAGCGTTGGCCG
ACAT

>a160_allele_B, aa 29
[CC]TCGGTGGACGTGACGACCCTGTTGAACGGCATCTCGACCGAGGACGGCCAGCTGGGAACGAACGCGAGGAGAAGTT
GTCGGTGCCTG**TGA**CGATCGTGAAAGGTTGCGTGTTGGGCTCCATCATAGTGACCGCGGTGTTCGGCAATCTGTTGGTGA
TGGTGTCGGTGATGCGGCACAGGAAGCTGCGGATCATCACCAATTATTTCGTGGTCTCTTTAGCGTTGGCCGACATGTTGG
TCGC

>b163_allele_A, aa 29
[CC]TCGGTGGACGTGACGACCCTGTTGAACGGCATCTCGACCGAGGACGGCCAGCTGGGAACGAACGCGAGGAGAAGTT
GTCGGTGCCTG**TGA**CGATCGTGAAAGGTTGCGTGTTGGGCTCCATCATAGTGACCGCGGTGTTCGGCAATCTGTTGGTGA
TGGTGTCGGTGATGCGGCACAGGAAGCTGCGGATCATCACCAATTATTTCGTGGTCTCTTTAGCGTTGGCCGACATGTTGG
TCGC

>b163_allele_B, aa 32
[CC]TCGGTGGACGTGACGACCCTGTTGAACGGCATCTCGACCGAGGACGGCCAGCTGGGAACGAACGCGAGCTACTCGA
GGAGAAGTTGTCGGTGCCTG**TGA**CGATCGTGAAAGGTTGCGTGTTGGGCTCCATCATAGTGACCGCGGTGTTCGGCAATC
TGTTGGTGATGGTGTCGGTGATGCGGCACAGGAAGCTGCGGATCATCACCAATTATTTCGTGGTCTCTTTAGCGTTGGCCG
ACAT

>a187_allele_A, aa 29
[CC]TCGGTGGACGTGACGACCCTGTTGAACGGCATCTCGACCGAGGACGGCCAGCTGGGAACGAACGCGAGGAGAAGTT
GTCGGTGCCTG**TGA**CGATCGTGAAAGGTTGCGTGTTGGGCTCCATCATAGTGACCGCGGTGTTCGGCAATCTGTTGGTGA
TGGTGTCGGTGATGCGGCACAGGAAGCTGCGGATCATCACCAATTATTTCGTGGTCTCTTTAGCGTTGGCCGACATGTTGG
TCGC

>a187_allele_B, aa 33
[CC]TCGGTGGACGTGACGACCCTGTTGAACGGCATCTCGACCGAGGACGGCCAGCTGGGAACGAACGCGAGCTACTCGA
GGAGGAGAAGTTGTCGGTGCCTG**TGA**CGATCGTGAAAGGTTGCGTGTTGGGCTCCATCATAGTGACCGCGGTGTTCGGCA
ATCTGTTGGTGATGGTGTCGGTGATGCGGCACAGGAAGCTGCGGATCATCACCAATTATTTCGTGGTCTCTTTAGCGTTGG
CCGA

>b158_allele_A, aa 23
[CC]TCGGTGGACGTGACGACCCTGTTGAACGGCATCTCGACCGAGGACGGCCAGCTGGGAACGAACGCGAGC**TAG**CGAG
GAGAAGTTGTCGGTGCCTGTGACGATCGTGAAAGGTTGCGTGTTGGGCTCCATCATAGTGACCGCGGTGTTCGGCAATCT
GTTGGTGATGGTGTCGGTGATGCGGCACAGGAAGCTGCGGATCATCACCAATTATTTCGTGGTCTCTTTAGCGTTGGCCGA
CATG

>b158_allele_B, aa 32
[CC]TCGGTGGACGTGACGACCCTGTTGAACGGCATCTCGACCGAGGACGGCCAGCTGGGAACGAACGCGAGCTACTCGA
GGAGAAGTTGTCGGTGCCTG**TGA**CGATCGTGAAAGGTTGCGTGTTGGGCTCCATCATAGTGACCGCGGTGTTCGGCAATC
TGTTGGTGATGGTGTCGGTGATGCGGCACAGGAAGCTGCGGATCATCACCAATTATTTCGTGGTCTCTTTAGCGTTGGCCG
ACAT

>a172_allele_A, aa 23
[CC]TCGGTGGACGTGACGACCCTGTTGAACGGCATCTCGACCGAGGACGGCCAGCTGGGAACGAACGCGAGC**TAG**CGAG
GAGAAGTTGTCGGTGCCTGTGACGATCGTGAAAGGTTGCGTGTTGGGCTCCATCATAGTGACCGCGGTGTTCGGCAATCT
GTTGGTGATGGTGTCGGTGATGCGGCACAGGAAGCTGCGGATCATCACCAATTATTTCGTGGTCTCTTTAGCGTTGGCCGA
CATG

>a172_allele_B, aa 32
[CC]TCGGTGGACGTGACGACCCTGTTGAACGGCATCTCGACCGAGGACGGCCAGCTGGGAACGAACGCGAGCTACTCGA
GGAGAAGTTGTCGGTGCCTG**TGA**CGATCGTGAAAGGTTGCGTGTTGGGCTCCATCATAGTGACCGCGGTGTTCGGCAATC
TGTTGGTGATGGTGTCGGTGATGCGGCACAGGAAGCTGCGGATCATCACCAATTATTTCGTGGTCTCTTTAGCGTTGGCCG
ACAT

>b132_allele_A, aa 32
[CC]TCGGTGGACGTGACGACCCTGTTGAACGGCATCTCGACCGAGGACGGCCAGCTGGGAACGAACGCGAGCTACTCGA
GGAGAAGTTGTCGGTGCCTG**TGA**CGATCGTGAAAGGTTGCGTGTTGGGCTCCATCATAGTGACCGCGGTGTTCGGCAATC

TGTTGGTGATGGTGTCGGTGATGCGGCACAGGAAGCTGCGGATCATCACCAATTATTTCGTGGTCTCTTTAGCGTTGGCCG
ACAT

>b132_allele_B, aa 32
[CC]TCGGTGGACGTGACGACCCTGTTGAACGGCATCTCGACCGAGGACGGCCAGCTGGGAACGAACGCGAGCTACGCGA
GGAGAAGTTGTCGGTGCCTG**TGA**CGATCGTGAAAGGTTGCGTGTTGGGCTCCATCATAGTGACCGCGGTGTTCGGCAATC
TGTTGGTGATGGTGTCGGTGATGCGGCACAGGAAGCTGCGGATCATCACCAATTATTTCGTGGTCTCTTTAGCGTTGGCCG
ACAT

>a259_allele_A, aa 32
[CC]TCGGTGGACGTGACGACCCTGTTGAACGGCATCTCGACCGAGGACGGCCAGCTGGGAACGAACGCGAGCTACGCGA
GGAGAAGTTGTCGGTGCCTG**TGA**CGATCGTGAAAGGTTGCGTGTTGGGCTCCATCATAGTGACCGCGGTGTTCGGCAATC
TGTTGGTGATGGTGTCGGTGATGCGGCACAGGAAGCTGCGGATCATCACCAATTATTTCGTGGTCTCTTTAGCGTTGGCCG
ACAT

>a259_allele_B, aa 32
[CC]TCGGTGGACGTGACGACCCTGTTGAACGGCATCTCGACCGAGGACGGCCAGCTGGGAACGAACGCGAGCTACTCGA
GGAGAAGTTGTCGGTGCCTG**TGA**CGATCGTGAAAGGTTGCGTGTTGGGCTCCATCATAGTGACCGCGGTGTTCGGCAATC
TGTTGGTGATGGTGTCGGTGATGCGGCACAGGAAGCTGCGGATCATCACCAATTATTTCGTGGTCTCTTTAGCGTTGGCCG
ACAT

>b190_allele_A, aa 32
[CC]TCGGTGGACGTGACGACCCTGTTGAACGGCATCTCGACCGAGGACGGCCAGCTGGGAACGAACGCGAGCTACGCGA
GGAGAAGTTGTCGGTGCCTG**TGA**CGATCGTGAAAGGTTGCGTGTTGGGCTCCATCATAGTGACCGCGGTGTTCGGCAATC
TGTTGGTGATGGTGTCGGTGATGCGGCACAGGAAGCTGCGGATCATCACCAATTATTTCGTGGTCTCTTTAGCGTTGGCCG
ACAT

>b190_allele_B, aa 32
[CC]TCGGTGGACGTGACGACCCTGTTGAACGGCATCTCGACCGAGGACGGCCAGCTGGGAACGAACGCGAGCTACTCGA
GGAGAAGTTGTCGGTGCCTG**TGA**CGATCGTGAAAGGTTGCGTGTTGGGCTCCATCATAGTGACCGCGGTGTTCGGCAATC
TGTTGGTGATGGTGTCGGTGATGCGGCACAGGAAGCTGCGGATCATCACCAATTATTTCGTGGTCTCTTTAGCGTTGGCCG
ACAT

>a155_allele_A, aa 32
[CC]TCGGTGGACGTGACGACCCTGTTGAACGGCATCTCGACCGAGGACGGCCAGCTGGGAACGAACGCGAGCTACGCGA
GGAGAAGTTGTCGGTGCCTG**TGA**CGATCGTGAAAGGTTGCGTGTTGGGCTCCATCATAGTGACCGCGGTGTTCGGCAATC
TGTTGGTGATGGTGTCGGTGATGCGGCACAGGAAGCTGCGGATCATCACCAATTATTTCGTGGTCTCTTTAGCGTTGGCCG
ACAT

>a155_allele_B, aa 32
[CC]TCGGTGGACGTGACGACCCTGTTGAACGGCATCTCGACCGAGGACGGCCAGCTGGGAACGAACGCGAGCTACTCGA
GGAGAAGTTGTCGGTGCCTG**TGA**CGATCGTGAAAGGTTGCGTGTTGGGCTCCATCATAGTGACCGCGGTGTTCGGCAATC
TGTTGGTGATGGTGTCGGTGATGCGGCACAGGAAGCTGCGGATCATCACCAATTATTTCGTGGTCTCTTTAGCGTTGGCCG
ACAT

>b234, aa 32
[CC]TCGGTGGACGTGACGACCCTGTTGAACGGCATCTCGACCGAGGACGGCCAGCTGGGAACGAACGCGAGCTACTCGA
GGAGAAGTTGTCGGTGCCTG**TGA**CGATCGTGAAAGGTTGCGTGTTGGGCTCCATCATAGTGACCGCGGTGTTCGGCAATC
TGTTGGTGATGGTGTCGGTGATGCGGCACAGGAAGCTGCGGATCATCACCAATTATTTCGTGGTCTCTTTAGCGTTGGCCG
ACAT

>a125, aa 32
[CC]TCGGTGGACGTGACGACCCTGTTGAACGGCATCTCGACCGAGGACGGCCAGCTGGGAACGAACGCGAGCTACTCGA
GGAGAAGTTGTCGGTGCCTG**TGA**CGATCGTGAAAGGTTGCGTGTTGGGCTCCATCATAGTGACCGCGGTGTTCGGCAATC
TGTTGGTGATGGTGTCGGTGATGCGGCACAGGAAGCTGCGGATCATCACCAATTATTTCGTGGTCTCTTTAGCGTTGGCCG
ACAT

>b250, aa 32
[CC]TCGGTGGACGTGACGACCCTGTTGAACGGCATCTCGACCGAGGACGGCCAGCTGGGAACGAACGCGAGCTACTCGA
GGAGAAGTTGTCGGTGCCTG**TGA**CGATCGTGAAAGGTTGCGTGTTGGGCTCCATCATAGTGACCGCGGTGTTCGGCAATC
TGTTGGTGATGGTGTCGGTGATGCGGCACAGGAAGCTGCGGATCATCACCAATTATTTCGTGGTCTCTTTAGCGTTGGCCG
ACAT

>a123, aa 32
[CC]TCGGTGGACGTGACGACCCTGTTGAACGGCATCTCGACCGAGGACGGCCAGCTGGGAACGAACGCGAGCTACTCGA
GGAGAAGTTGTCGGTGCCTG**TGA**CGATCGTGAAAGGTTGCGTGTTGGGCTCCATCATAGTGACCGCGGTGTTCGGCAATC
TGTTGGTGATGGTGTCGGTGATGCGGCACAGGAAGCTGCGGATCATCACCAATTATTTCGTGGTCTCTTTAGCGTTGGCCG
ACAT

>b238, aa 32
[CC]TCGGTGGACGTGACGACCCTGTTGAACGGCATCTCGACCGAGGACGGCCAGCTGGGAACGAACGCGAGCTACTCGA
GGAGAAGTTGTCGGTGCCTG**TGA**CGATCGTGAAAGGTTGCGTGTTGGGCTCCATCATAGTGACCGCGGTGTTCGGCAATC
TGTTGGTGATGGTGTCGGTGATGCGGCACAGGAAGCTGCGGATCATCACCAATTATTTCGTGGTCTCTTTAGCGTTGGCCG
ACAT

>a102, aa 32
[CC]TCGGTGGACGTGACGACCCTGTTGAACGGCATCTCGACCGAGGACGGCCAGCTGGGAACGAACGCGAGCTACTCGA
GGAGAAGTTGTCGGTGCCTG**TGA**CGATCGTGAAAGGTTGCGTGTTGGGCTCCATCATAGTGACCGCGGTGTTCGGCAATC
TGTTGGTGATGGTGTCGGTGATGCGGCACAGGAAGCTGCGGATCATCACCAATTATTTCGTGGTCTCTTTAGCGTTGGCCG
ACAT

>b92_allele_A, aa 32
[CC]TCGGTGGACGTGACGACCCTGTTGAACGGCATCTCGACCGAGGACGGCCAGCTGGGAACGAACGCGAGCTACTCGA
GGAGAAGTTGTCGGTGCCTG**TGA**CGATCGTGAAAGGTTGCGTGTTGGGCTCCATCATAGTGACCGCGGTGTTCGGCAATC
TGTTGGTGATGGTGTCGGTGATGCGGCACAGGAAGCTGCGGATCATCACCAATTATTTCGTGGTCTCTTTAGCGTTGGCCG
ACAT

>b92_allele_B, aa 32
[CC]TCGGTGGACGTGACGACCCTGTTGAACGGCATCTCGACCGAGGACGGCCAGCTGGGAACGAACGCGAGCTACGCGA
GGAGAAGTTGTCGGTGCCTG**TGA**CGATCGTGAAAGGTTGCGTGTTGGGCTCCATCATAGTGACCGCGGTGTTCGGCAATC
TGTTGGTGATGGTGTCGGTGATGCGGCACAGGAAGCTGCGGATCATCACCAATTATTTCGTGGTCTCTTTAGCGTTGGCCG
ACAT

>a244_allele_A, aa 29
[CC]TCGGTGGACGTGACGACCCTGTTGAACGGCATCTCGACCGAGGACGGCCAGCTGGGAACGAACGCGAGGAGAAGTT
GTCGGTGCCTG**TGA**CGATCGTGAAAGGTTGCGTGTTGGGCTCCATCATAGTGACCGCGGTGTTCGGCAATCTGTTGGTGA
TGGTGTCGGTGATGCGGCACAGGAAGCTGCGGATCATCACCAATTATTTCGTGGTCTCTTTAGCGTTGGCCGACATGTTGG
TCGC

>a244_allele_B, aa 35
[CC]TCGGTGGACGTGACGACCCTGTTGAACGGCATCTCGACCGAGGACGGCCAGCTGGGAACGAACGCGAGCTACTCGA
TGAACGCGAGGAGAAGTTGTCGGTGCCTG**TGA**CGATCGTGAAAGGTTGCGTGTTGGGCTCCATCATAGTGACCGCGGTGT
TCGGCAATCTGTTGGTGATGGTGTCGGTGATGCGGCACAGGAAGCTGCGGATCATCACCAATTATTTCGTGGTCTCTTTAG
CGTT

>b153, aa 32
[CC]TCGGTGGACGTGACGACCCTGTTGAACGGCATCTCGACCGAGGACGGCCAGCTGGGAACGAACGCGAGCTACTCGA
GGAGAAGTTGTCGGTGCCTG**TGA**CGATCGTGAAAGGTTGCGTGTTGGGCTCCATCATAGTGACCGCGGTGTTCGGCAATC
TGTTGGTGATGGTGTCGGTGATGCGGCACAGGAAGCTGCGGATCATCACCAATTATTTCGTGGTCTCTTTAGCGTTGGCCG
ACAT

>a248_allele_A, aa 32
[CC]TCGGTGGACGTGACGACCCTGTTGAACGGCATCTCGACCGAGGACGGCCAGCTGGGAACGAACGCGAGCTACTCGA
GGAGAAGTTGTCGGTGCCTG**TGA**CGATCGTGAAAGGTTGCGTGTTGGGCTCCATCATAGTGACCGCGGTGTTCGGCAATC
TGTTGGTGATGGTGTCGGTGATGCGGCACAGGAAGCTGCGGATCATCACCAATTATTTCGTGGTCTCTTTAGCGTTGGCCG
ACAT

>a248_allele_B, aa 32
[CC]TCGGTGGACGTGACGACCCTGTTGAACGGCATCTCGACCGAGGACGGCCAGCTGGGAACGAACGCGAGCTACTCGA
GGAGGAGTTGTCGGTGCCTG**TGA**CGATCGTGAAAGGTTGCGTGTTGGGCTCCATCATAGTGACCGCGGTGTTCGGCAATC
TGTTGGTGATGGTGTCGGTGATGCGGCACAGGAAGCTGCGGATCATCACCAATTATTTCGTGGTCTCTTTAGCGTTGGCCG
ACAT

>b161_allele_A, aa 32
[CC]TCGGTGGACGTGACGACCCTGTTGAACGGCATCTCGACCGAGGACGGCCAGCTGGGAACGAACGCGAGCTACTCGA
GGAGAAGTTGTCGGTGCCTG**TGA**CGATCGTGAAAGGTTGCGTGTTGGGCTCCATCATAGTGACCGCGGTGTTCGGCAATC
TGTTGGTGATGGTGTCGGTGATGCGGCACAGGAAGCTGCGGATCATCACCAATTATTTCGTGGTCTCTTTAGCGTTGGCCG
ACAT

>b161_allele_B, aa 32
[CC]TCGGTGGACGTGACGACCCTGTTGAACGGCATCTCGACCGAGGACGGCCAGCTGGGAACGAACGCGAGCTACTCGA
GGAGGAGTTGTCGGTGCCTG**TGA**CGATCGTGAAAGGTTGCGTGTTGGGCTCCATCATAGTGACCGCGGTGTTCGGCAATC
TGTTGGTGATGGTGTCGGTGATGCGGCACAGGAAGCTGCGGATCATCACCAATTATTTCGTGGTCTCTTTAGCGTTGGCCG
ACAT

>a183_allele_A, aa 32
[CC]TCGGTGGACGTGACGACCCTGTTGAACGGCATCTCGACCGAGGACGGCCAGCTGGGAACGAACGCGAGCTACGCGA
GGAGAAGTTGTCGGTGCCTG**TGA**CGATCGTGAAAGGTTGCGTGTTGGGCTCCATCATAGTGACCGCGGTGTTCGGCAATC
TGTTGGTGATGGTGTCGGTGATGCGGCACAGGAAGCTGCGGATCATCACCAATTATTTCGTGGTCTCTTTAGCGTTGGCCG
ACAT

>a183_allele_B, aa 80 nonsense
[CC]TCGGTGGACGTGACGACCCTGTTGAACGGCATCTCGACCGAGGACGGCCAGCTGGGAACGAACGCGAGCTACTCGC
GAGGAGAAGTTGTCGGTGCCTGTGACGATCGTGAAAGGTTGCGTGTTGGGCTCCATCATAGTGACCGCGGTGTTCGGCAA
TCTGTTGGTGATGGTGTCGGTGATGCGGCACAGGAAGCTGCGGATCATCACCAATTATTTCGTGGTCTCTTTAGCGTTGGC
CGAC

>b107_allele_A, aa 32
[CC]TCGGTGGACGTGACGACCCTGTTGAACGGCATCTCGACCGAGGACGGCCAGCTGGGAACGAACGCGAGCTACTCGA

GGAGAAGTTGTCGGTGCCTG**TGA**CGATCGTGAAAGGTTGCGTGTTGGGCTCCATCATAGTGACCGCGGTGTTCGGCAATC
TGTTGGTGATGGTGTCGGTGATGCGGCACAGGAAGCTGCGGATCATCACCAATTATTTCGTGGTCTCTTTAGCGTTGGCCG
ACAT

>a98, aa 32
[CC]TCGGTGGACGTGACGACCCTGTTGAACGGCATCTCGACCGAGGACGGCCAGCTGGGAACGAACGCGAGCTACTCGA
GGAGAAGTTGTCGGTGCCTG**TGA**CGATCGTGAAAGGTTGCGTGTTGGGCTCCATCATAGTGACCGCGGTGTTCGGCAATC
TGTTGGTGATGGTGTCGGTGATGCGGCACAGGAAGCTGCGGATCATCACCAATTATTTCGTGGTCTCTTTAGCGTTGGCCG
ACAT

>b93_allele_A, aa 32
[CC]TCGGTGGACGTGACGACCCTGTTGAACGGCATCTCGACCGAGGACGGCCAGCTGGGAACGAACGCGAGCTACTCGA
GGAGAAGTTGTCGGTGCCTG**TGA**CGATCGTGAAAGGTTGCGTGTTGGGCTCCATCATAGTGACCGCGGTGTTCGGCAATC
TGTTGGTGATGGTGTCGGTGATGCGGCACAGGAAGCTGCGGATCATCACCAATTATTTCGTGGTCTCTTTAGCGTTGGCCG
ACAT

>b93_allele_B, aa 80 nonsense
[CC]TCGGTGGACGTGACGACCCTGTTGAACGGCATCTCGACCGAGGACGGCCAGCTGGGAACGAACGCGAGCTACTCGC
GAGGAGAAGTTGTCGGTGCCTGTGACGATCGTGAAAGGTTGCGTGTTGGGCTCCATCATAGTGACCGCGGTGTTCGGCAA
TCTGTTGGTGATGGTGTCGGTGATGCGGCACAGGAAGCTGCGGATCATCACCAATTATTTCGTGGTCTCTTTAGCGTTGGC
CGAC

>a242_allele_A, aa 80 nonsense
[CC]TCGGTGGACGTGACGACCCTGTTGAACGGCATCTCGACCGAGGACGGCCAGCTGGGAACGAACGCGAGCTACTCGC
GAGGAGAAGTTGTCGGTGCCTGTGACGATCGTGAAAGGTTGCGTGTTGGGCTCCATCATAGTGACCGCGGTGTTCGGCAA
TCTGTTGGTGATGGTGTCGGTGATGCGGCACAGGAAGCTGCGGATCATCACCAATTATTTCGTGGTCTCTTTAGCGTTGGC
CGAC

>a242_allele_B, aa 32
[CC]TCGGTGGACGTGACGACCCTGTTGAACGGCATCTCGACCGAGGACGGCCAGCTGGGAACGAACGCGAGCTACTCGA
GGAGAAGTTGTCGGTGCCTG**TGA**CGATCGTGAAAGGTTGCGTGTTGGGCTCCATCATAGTGACCGCGGTGTTCGGCAATC
TGTTGGTGATGGTGTCGGTGATGCGGCACAGGAAGCTGCGGATCATCACCAATTATTTCGTGGTCTCTCTAGCGTTGGCCG
ACAT

>b169_allele_A, aa 80 nonsense
[CC]TCGGTGGACGTGACGACCCTGTTGAACGGCATCTCGACCGAGGACGGCCAGCTGGGAACGAACGCGAGCTACTCGC
GAGGAGAAGTTGTCGGTGCCTGTGACGATCGTGAAAGGTTGCGTGTTGGGCTCCATCATAGTGACCGCGGTGTTCGGCAA
TCTGTTGGTGATGGTGTCGGTGATGCGGCACAGGAAGCTGCGGATCATCACCAATTATTTCGTGGTCTCTTTAGCGTTGGC
CGAC

>b169_allele_B, aa 32
[CC]TCGGTGGACGTGACGACCCTGTTGAACGGCATCTCGACCGAGGACGGCCAGCTGGGAACGAACGCGAGCTACTCGA
GGAGAAGTTGTCGGTGCCTG**TGA**CGATCGTGAAAGGTTGCGTGTTGGGCTCCATCATAGTGACCGCGGTGTTCGGCAATC
TGTTGGTGATGGTGTCGGTGATGCGGCACAGGAAGCTGCGGATCATCACCAATTATTTCGTGGTCTCTCTAGCGTTGGCCG
ACAT

>a232, aa 80 nonsense
[CC]TCGGTGGACGTGACGACCCTGTTGAACGGCATCTCGACCGAGGACGGCCAGCTGGGAACGAACGCGAGCTACTCGC
GAGGAGAAGTTGTCGGTGCCTGTGACGATCGTGAAAGGTTGCGTGTTGGGCTCCATCATAGTGACCGCGGTGTTCGGCAA
TCTGTTGGTGATGGTGTCGGTGATGCGGCACAGGAAGCTGCGGATCATCACCAATTATTTCGTGGTCTCTTTAGCGTTGGC
CGAC

>b218, aa 80 nonsense
[CC]TCGGTGGACGTGACGACCCTGTTGAACGGCATCTCGACCGAGGACGGCCAGCTGGGAACGAACGCGAGCTACTCGC
GAGGAGAAGTTGTCGGTGCCTGTGACGATCGTGAAAGGTTGCGTGTTGGGCTCCATCATAGTGACCGCGGTGTTCGGCAA
TCTGTTGGTGATGGTGTCGGTGATGCGGCACAGGAAGCTGCGGATCATCACCAATTATTTCGTGGTCTCTTTAGCGTTGGC
CGAC

>a184, aa 80 nonsense
[CC]TCGGTGGACGTGACGACCCTGTTGAACGGCATCTCGACCGAGGACGGCCAGCTGGGAACGAACGCGAGCTACTCGC
GAGGAGAAGTTGTCGGTGCCTGTGACGATCGTGAAAGGTTGCGTGTTGGGCTCCATCATAGTGACCGCGGTGTTCGGCAA
TCTGTTGGTGATGGTGTCGGTGATGCGGCACAGGAAGCTGCGGATCATCACCAATTATTTCGTGGTCTCTTTAGCGTTGGC
CGAC

>b237, aa 33
[CC]TCGGTGGACGTGACGACCCTGTTGAACGGCATCTCGACCGAGGACGGCCAGCTGGGAACGAACGCGAGCTACTTCG
CGAGGAGAAGTTGTCGGTGCCTG**TGA**CGATCGTGAAAGGTTGCGTGTTGGGCTCCATCATAGTGACCGCGGTGTTCGGCA
ATCTGTTGGTGATGGTGTCGGTGATGCGGCACAGGAAGCTGCGGATCATCACCAATTATTTCGTGGTCTCTTTAGCGTTGG
CCGA

>a201_allele_A, aa 80 nonsense
[CC]TCGGTGGACGTGACGACCCTGTTGAACGGCATCTCGACCGAGGACGGCCAGCGAGGAGAAGTTGTCGGTGCCTGTG
ACGATCGTGAAAGGTTGCGTGTTGGGCTCCATCATAGTGACCGCGGTGTTCGGCAATCTGTTGGTGATGGTGTCGGTGAT

GCGGCACAGGAAGCTGCGGATCATCACCAATTATTTCGTGGTCTCTTTAGCGTTGGCCGACATGTTGGTCGCCATGTTCGC
CATG

>a201_allele_B, aa 30
[CC]TCGGTGGACGTGACGACCCTGTTGAACGGCATCTCGACCGAGGACGGCCAGCTGGGAACGAACGCGAGCTGGAGAA
GTTGTCGGTGCCTG**TGA**CGATCGTGAAAGGTTGCGTGTTGGGCTCCATCATAGTGACCGCGGTGTTCGGCAATCTGTTGG
TGATGGTGTCGGTGATGCGGCACAGGAAGCTGCGGATCATCACCAATTATTTCGTGGTCTCTTTAGCGTTGGCCGACATGT
TGGT

>b191, aa 29
[CC]TCGGTGGACGTGACGACCCTGTTGAACGGCATCTCGACCGAGGACGGCCAGCTGGGAACGAACGCGAGGAGAAGTT
GTCGGTGCCTG**TGA**CGATCGTGAAAGGTTGCGTGTTGGGCTCCATCATAGTGACCGCGGTGTTCGGCAATCTGTTGGTGA
TGGTGTCGGTGATGCGGCACAGGAAGCTGCGGATCATCACCAATTATTTCGTGGTCTCTTTAGCGTTGGCCGACATGTTGG
TCGC
